# Engineering CAR-T cells to remodel the mucin-rich cancer cell glycocalyx

**DOI:** 10.64898/2026.09.25.754490

**Authors:** Sangwoo Park, Cassidy E. Ho, Alexandra N. Wolff, Yueyang Fan, Amanda A. Bouffard, Justin Paek, Adele Mucci, Trisha R. Berger, Matthew J. Paszek, Marcela V. Maus

**Author notes:** Corresponding author. &.

## Abstract

The dense glycocalyx of cancer cells can restrict immune-cell access to surface antigens and limit CAR-T cell activity. Here, we show that mucin density and epitope position determine how glycocalyx remodeling affects CAR-T cell recognition and killing. We identify KLK5 as a human protease that cleaves tumor-associated mucins, increases access to membrane-proximal antigens, and enhances CAR-T cell function. We then engineer CAR-T cells to display or secrete KLK5, enabling remodeling of the tumor glycocalyx during antigen recognition. KLK5-engineered CAR-T cells improved tumor control across multiple xenograft models, and KLK5-secreting MUC17 CAR-T cells produced the strongest in vivo benefit, prolonging survival compared with conventional MUC17 CAR-T cells. These findings show that CAR-T cells can be engineered to breach the mucin-rich glycocalyx while preserving accessible target epitopes.

## Main Text

Chimeric antigen receptor (CAR)-T cell therapies have produced remarkable clinical responses in hematological malignancies(*1*). However, their efficacy in solid tumors remains limited by heterogeneous antigen expression, poor trafficking and infiltration, immunosuppressive signals, and progressive loss of T-cell function within the tumor microenvironment. Several engineering approaches have been developed to address these barriers, including multi-antigen targeting CARs, synthetic receptor systems, and genetic perturbations that improve T-cell function(*2–5*). Most of these approaches modify antigen recognition by the CAR, downstream signaling, or the functional state of the T cell. However, antigen recognition by the CAR also requires close physical contact with the tumor cell surface, which is not directly addressed by these strategies.

One regulator of these cell surface interactions is the cellular glycocalyx, which is composed of glycoproteins, glycolipids, and proteoglycans that extend from the plasma membrane. Changes in glycocalyx composition on cancer cells can alter receptor organization, adhesion, and signaling(*6*). Likewise, long glycocalyx biopolymers also generate steric and entropic forces that increase with molecular size and surface density(*7*). Major structural components of the cancer glycocalyx are transmembrane mucins, which contain extended extracellular domains that are densely modified with O-linked glycans. Mucins are frequently overexpressed or aberrantly glycosylated in epithelial cancers and can contribute to tumor progression and immune evasion through both biochemical and physical mechanisms(*8*, *9*). We previously showed that mucin density, glycosylation, and backbone length regulate the nanoscale thickness of the cancer cell glycocalyx, with even ∼10 nm differences in thickness altering susceptibility to cytotoxic immune cell attack(*10*, *11*). Disruption of tumor cell *N*-glycosylation has also been shown to improve immune synapse formation and CAR-T cell activity, further supporting a role for the glycocalyx in limiting engineered immune cell function(*12*).

Mucins present an additional problem when they are used as the CAR target antigen. Tumor-associated mucins can provide abundant CAR-binding sites. For example, Tn-MUC1 is a tumor-associated mucin glycoform that can be directly targeted by CAR-T cells(*13*). However, high mucin surface density can also increase the physical barrier to receptor engagement. Therefore, increasing the expression of a mucin antigen may have opposing effects on CAR T cell activation by increasing the number of available CAR-binding sites while also increasing steric resistance within the glycocalyx(*7*, *10*). CAR-T cell activation can also depend on the position of the targeted epitope. Membrane-proximal epitopes can promote stronger receptor activation than membrane-distal epitopes by favoring close membrane apposition and exclusion of the large phosphatase CD45 from the receptor-ligand interface(*14*). Thus, both mucin density and epitope position may affect CAR-T cell recognition, but how these two properties interact during mucin-directed CAR-T cell attack remains unclear.

One approach to reduce physical barriers encountered by engineered T cells is through enzymatic remodeling. For example, CAR-T cells expressing heparanase can degrade extracellular-matrix heparan sulfate, which increases tumor infiltration and improves antitumor activity(*15*). The glycocalyx differs from the surrounding extracellular matrix in that it is positioned directly at the receptor-ligand interface. We previously showed that equipping cytotoxic immune cells with StcE, a mucin-selective bacterial protease, reduces the mucin barrier and enhances cytotoxicity(*10*). Tumor-targeted StcE has subsequently been used to remove cancer-associated mucins while reducing systemic exposure to mucinase activity(*16*). Similarly, enzymatic editing of tumor-cell glycans with antibody-sialidase conjugates similarly show that enzymatic editing of tumor-cell glycans can enhance antitumor immunity(*17*). However, these approaches rely on exogenously administered enzymes, including enzymes of microbial origin. Therefore, we asked whether a human protease could be genetically delivered by CAR-T cells to remodel the tumor cell glycocalyx during tumor cell engagement.

Here, we investigate the impact of mucin density and epitope position on CAR-T cell recognition of cancer cells and develop a human protease-based approach to remodel the tumor-cell glycocalyx. We show that increasing Tn-MUC1 density initially enhances CAR-T cell killing but reduces cytotoxicity at higher surface densities. In contrast, proteolytic removal of the distal mucin layer increases accessibility to membrane-proximal MUC1 and MUC17 epitopes and enhances MUC17 CAR-T cell avidity and killing. We identify kallikrein-related peptidase 5 (KLK5) as a human protease that cleaves several glycocalyx proteins and reduces glycocalyx thickness. We further engineer CAR-T cells to either express KLK5 using an ADAM10-based membrane-tethered enzyme receptor or secrete soluble KLK5. Both approaches increase CAR-T cell activity in gastric and pancreatic tumor models, although the effect depends on whether the targeted epitope is preserved following proteolysis. Overall, our results show that the physical organization of the tumor cell glycocalyx can influence CAR-T cell recognition and can be modified directly by engineered T cells.

## Results

### Mucin density and epitope position regulate CAR-T cell recognition and cytotoxicity

Cell-surface mucins form an extended glycocalyx that can restrict access to molecules positioned closer to the plasma membrane. We previously showed that higher mucin expression increases the nanoscale thickness of the glycocalyx and reduces physical interactions between cytotoxic immune cells and target cells(*10*). Therefore, we asked whether mucin cleavage would differentially alter target recognition, depending on the position of the antibody-binding epitope (**Fig. 1A,B**). To test this possibility over a controlled range of mucin expression, we utilized a doxycycline-inducible MUC1 system in cells with a with *C1GALT1* knockout, which produces the truncated Tn glycoform of MUC1 (Tn-MUC1)(*10*) (**Fig. 1C,D**). Tn-MUC1 is a tumor-associated glycoform that has previously been targeted using CAR-T cells(*13*). In this system, increasing doxycycline progressively increased cell-surface Tn-MUC1. We compared recognition of the readily available MUC1 tandem-repeat, which is a more membrane-distal region, with recognition of the membrane-proximal MUC1-C domain before and after treatment with the mucin-selective protease StcE. StcE treatment did not increase recognition of the tandem-repeat epitope, and modestly reduced its recognition at higher MUC1 expression levels, likely due to shedding of the antigen-binding domains. In contrast, binding of the membrane-proximal MUC1-C antibody increased by approximately 1.7-fold following StcE treatment (**Fig. 1E**). Thus, removal of the extended mucin ectodomain had different effects on antibody recognition depending on the position of the target epitope.

**Fig 1.**
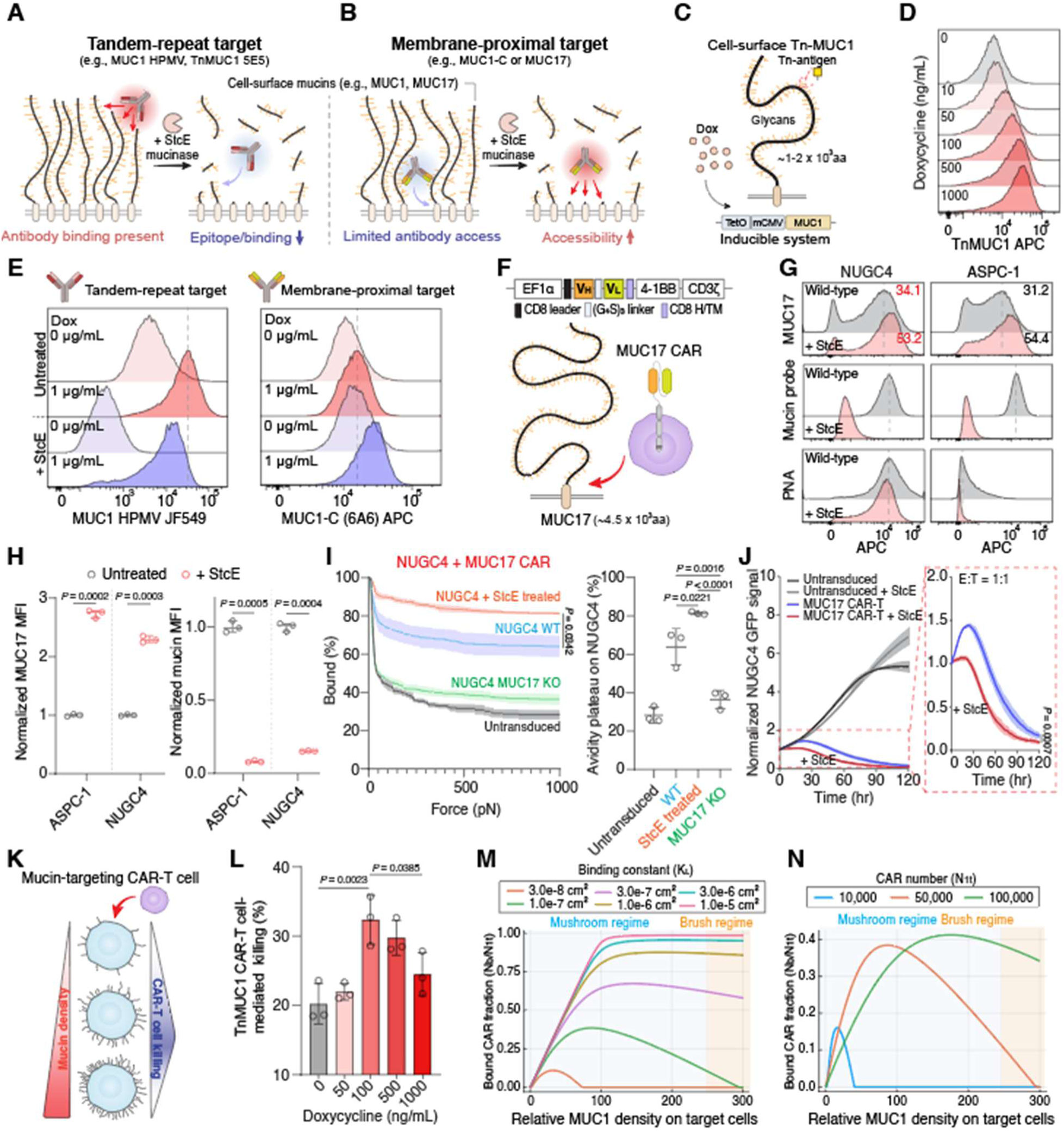
Mucin density regulates antibody recognition, CAR-T cell avidity and cytotoxicity. (**A** and **B**) Schematic illustrating antibody recognition of epitopes located within the tandem-repeat region of mucins (**A**) or at membrane-proximal sites (**B**). StcE-mediated cleavage of cell-surface mucins differentially affects antibody recognition depending on epitope location. (**C**) Schematic of the doxycycline-inducible system used to control Tn-MUC1 expression through *C1GALT1* knockout. (**D**) Flow cytometry analysis of Tn-MUC1 expression at the indicated concentrations of doxycycline. (**E**) Flow cytometry analysis of MUC1 expression using antibodies targeting the tandem-repeat region (MUC1 HMPV; left) or a membrane-proximal MUC1-C epitope (6A6; right) at the indicated doxycycline concentrations before and after treatment with 100 nM StcE. (**F**) Schematic of MUC17 CAR-T cells recognizing a membrane-proximal epitope of MUC17. (**G**) Flow cytometry analysis of MUC17, mucin-probe and PNA binding to NUGC4 and ASPC-1 cells before and after treatment with 100 nM StcE. (**H**) Quantification of normalized MUC17 and mucin-probe mean fluorescence intensity (MFI) from Fig. 1G. Results are mean ± s.d. of n = 3 independent measurements. (**I**) Measurement of cell avidity between untreated or StcE-treated NUGC4 cells or MUC17-knockout NUGC4 cells and either MUC17 CAR-T or untransduced T cells. The percentage of T cells remaining bound to target cells is shown as the applied acoustic force is increased from 0 to 1,000 pN (left). The percentage of CAR-T cells remaining bound at the plateau of an avidity at 1,000 pN is also quantified (right). Results are mean ± s.e.m. for force-ramp measurements and mean ± s.d. for avidity-plateau measurements from n = 3 independent measurements. (**J**) Representative real-time cytotoxicity assay of MUC17 CAR-T cells against untreated or StcE-treated NUGC4 cells at 1:1 E:T ratio. Results are mean ± s.d. of n = 3 independent measurements. (**K**) Conceptual model illustrating the proposed relationship between mucin density and the activity of mucin-targeting CAR-T cells. Increasing mucin density initially increases target availability and CAR-T cell killing; above an optimal density, however, the mucin-rich glycocalyx acts as a steric barrier that limits CAR-T cell engagement and reduces cytotoxicity. (**L**) Luciferase-based cytotoxicity assay of Tn-MUC1-targeting CAR-T cells against cells with doxycycline-inducible Tn-MUC1 expression at the indicated doxycycline concentrations. (**M** and **N**) Theoretical predictions of the relationship between mucin density and MUC1 CAR-T cell activity as a function of CAR-antigen binding affinity (**M**) or CAR surface expression (**N**). In (**J**), statistical analysis was performed by two-way ANOVA with correction for multiple comparisons. In (**G**), (**I**), and (**L**), statistical analysis was performed by one-way ANOVA with Tukey’s post hoc tests.

A similar effect was observed with the endogenous mucin, MUC17, on cancer cells. MUC17 is a large membrane-tethered mucin expressed in subsets of gastrointestinal cancers and can be targeted using CAR-T cells(*18*). Given its extended extracellular domain and the membrane-proximal location of the CAR-targeted epitope, we used MUC17 as a proof-of-concept system to investigate how glycocalyx physical structure influences antigen accessibility and CAR-T cell recognition (**Fig. 1F**). StcE treatment reduced the overall cell-surface mucin levels but increased MUC17 antibody binding by approximately 2.7-fold in ASPC-1 pancreatic cancer cells and 2.3-fold in NUGC4 gastric cancer cells (**Fig. 1G,H, and fig. S1A**). Thus, reduction of the distal mucin layer increased accessibility to a membrane-proximal MUC17 epitope while also reducing the overall mucin-rich glycocalyx.

To determine whether increased MUC17 accessibility translated into stronger CAR-T cell interactions, we measured the interaction strength between MUC17 CAR-T cells and NUGC4 and GSU target cells using an acoustic force-based avidity assay (**Fig. 1I and fig. S1B,C**). As the applied force increased, StcE treatment increased the fraction of MUC17 CAR-T cells that remained bound to target cells, whereas binding to MUC17 KO cells or binding of untransduced T cells remained low. Consistent with the increased avidity, StcE-treated NUGC4 cells were more efficiently killed by MUC17 CAR-T cells (**Fig. 1J and fig. S1D**). These results indicate that mucin removal can improve CAR-T cell avidity and cytotoxicity when the targeted epitope is located close to the plasma membrane. By contrast, when a CAR directly recognizes an epitope within the mucin ectodomain itself, cell-surface mucin expression creates a competing effect: increasing expression raises antigen density but also increases the steric barrier surrounding the target. Using the inducible Tn-MUC1 system, we observed that increasing Tn-MUC1 expression initially enhanced CAR-T cell killing, from 20.2% without doxycycline to 32.4% at 100 ng/mL doxycycline. However, further increasing doxycycline to 1,000 ng/mL reduced cytotoxicity to 24.5%, despite the greater expression of target antigen (**Fig. 1L**). Thus, when the CAR-targeted epitope resides within the extended mucin ectodomain, increasing mucin expression simultaneously increases antigen expression and the physical barrier presented by the glycocalyx. At high mucin densities, this barrier became dominant, resulting in reduced CAR-T cell activity despite greater target expression.

To quantitatively examine this competition between antigen-dependent adhesion and the glycocalyx barrier, we developed a theoretical model of CAR-T-cell binding to tumor cells. Antigen density, CAR expression and CAR-antigen affinity can each contribute to CAR engagement(*19–22*). Thus, we adapted a thermodynamic cell-adhesion model developed by Bell and colleagues, in which receptor-ligand binding is determined by the equilibrium formation of cell-cell bonds and the energetic cost of bond extension. We extended this framework by adding a glycocalyx-repulsion term derived from a polymer-brush model, such that the total free energy included contributions from CAR-antigen binding and compression of the glycocalyx within the cell-cell contact area. The total free energy was minimized with respect to the number of CAR- antigen bonds, contact area and intermembrane separation to determine the equilibrium state for each set of binding and glycocalyx parameters. We first modeled MUC1-directed CAR binding (**Fig. 1M,N**) and subsequently examined MUC17-directed binding (**fig. S2**). In both cases, the CAR-binding site was represented as an ectodomain target without explicitly incorporating differences in epitope position along the mucin backbone. We then varied glycocalyx density together with CAR-antigen affinity or CAR expression to examine how these parameters shifted the balance between adhesion and repulsion. The model predicted a similar non-monotonic relationship between mucin density and CAR engagement. Increasing CAR-antigen affinity shifted the balance toward adhesion and partially reduced the inhibitory effect of increasing mucin density. Increasing CAR expressions produced a similar effect. However, at high mucin densities, the glycocalyx barrier reduced CAR-antigen binding even when CAR affinity or expression was increased. Thus, the model was consistent with our experimental observations and provided a framework for predicting how changes in CAR affinity, CAR expression, and glycocalyx density together determine receptor engagement. Overall, these results indicate that the antigen density experienced by a CAR-T cell depends on both the expression of the target and the physical environment in which it is presented.

## KLK5 remodels the mucin-rich cancer glycocalyx

StcE was effective at testing whether removal of the mucin layer could improve CAR-T cell engagement in vitro. However, StcE is a mucin-selective protease derived from enterohemorrhagic Escherichia coli and, when systemically administered in mice, StcE can cleave mucins in normal tissues and exhibits a relatively narrow tolerability window, motivating approaches that spatially restrict its proteolytic activity in vivo(*16*). Therefore, we sought to identify a human extracellular protease capable of remodeling the cancer-associated glycocalyx. This concept was supported by recent work showing that cathepsin K can degrade several components of the glycocalyx, further suggesting that human proteases could be used for this purpose(*23*).

To identify a human extracellular protease capable of remodeling the mucin-rich glycocalyx, we focused on kallikrein-related peptidases (KLKs), a family of extracellular serine proteases with well-characterized activity in epithelial tissues (**Fig. 2A**). We screened recombinant human KLK4, KLK5, KLK6, KLK7, and KLK13 for their ability to cleave MUC1 on NUGC4 cells (**Fig. 2B**). Western blot analysis showed the greatest MUC1 cleavage following KLK5 treatment, and we therefore selected KLK5 for further study. KLK5 also cleaved purified recombinant MUC17 and MUC16 (**Fig. 2C**), indicating activity across multiple large mucins and supporting its potential as a broader glycocalyx-remodeling protease.

**Fig. 2.**
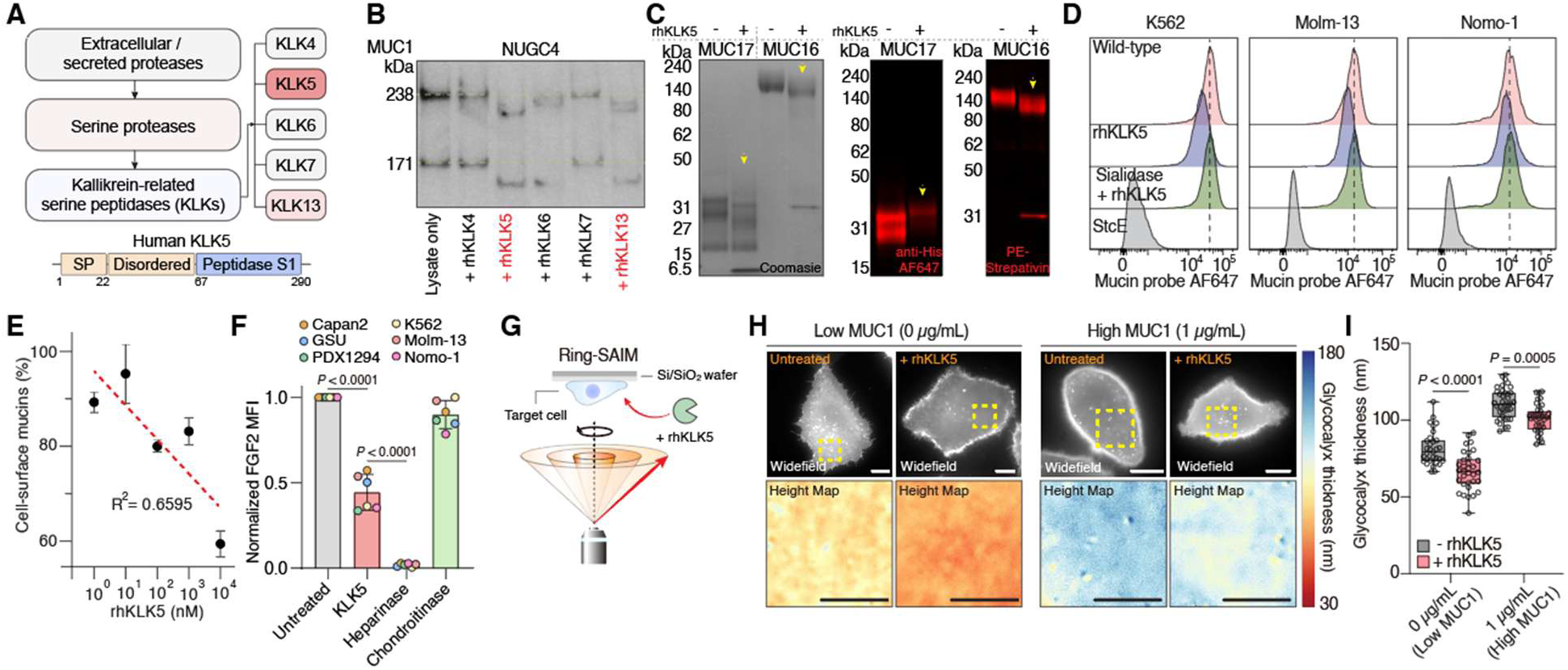
KLK5 is a mucin-modifying protease that remodels the glycocalyx and reduces glycocalyx thickness. (**A**) Schematic illustrating the selection of kallikrein-related serine peptidases (KLKs) for screening. (**B**) Western blot analysis of MUC1 in NUGC4 cells after treatment with recombinant human KLK4, KLK5, KLK6, KLK7 or KLK13. (**C**) Coomassie staining and western blot analysis of recombinant MUC17 and MUC16 following treatment with KLK5. (**D**) Flow cytometry analysis of cell-surface mucins using a mucin probe in K562, Molm-13 and Nomo-1 cells after treatment with recombinant human KLK5 (rhKLK5), sialidase followed by rhKLK5, or StcE. Cells were treated with the indicated enzymes for 2 h at 37 °C. (**E**) Cell-surface mucin levels on K562 cells after treatment with rhKLK5 at the indicated concentrations. The dashed line indicates a linear fit to the data. Results are mean ± s.d. of n = 3 independent measurements. (**F**) Normalized FGF2 binding on the indicated cell lines after treatment with KLK5, heparinase or chondroitinase for 2 h at 37 °C. (**G**) Schematic of scanning angle interference microscopy (SAIM) for measuring glycocalyx thickness before and after rhKLK5 treatment. (**H**) Representative wide-field image and corresponding glycocalyx-thickness map of live MCF10A cells expressing MUC1-GFP at the indicated doxycycline induction levels. Scale bars, 10 µm. (**I**) Quantification of glycocalyx thickness in MCF10A cells before and after treatment with rhKLK5. Boxes and whiskers indicate the first and third quartiles (boxes), median and range. Each condition includes a minimum of 31 cells from one representative experiment; n = 3 independent experiments. Statistical analysis in (**F)** and **(I**) were performed using one-way ANOVA with Tukey’s multiple-comparisons test.

KLK5 activity was not limited to mucins. CD43 is a heavily O-glycosylated and sialylated cell-surface protein recently shown to function as a glyco-immune barrier in acute myeloid leukemia(*24*). KLK5 treatment substantially cleaved recombinant CD43, whereas recombinant PSGL-1 and CD45, two other heavily glycosylated leukocyte surface proteins, were less affected under the same conditions (**fig. S3A**). Thus, KLK5 did not uniformly cleave highly glycosylated cell-surface proteins tested. To further examine KLK5 cleavage of MUC1, we analyzed KLK5-generated MUC1 products by mass spectrometry. Prominent cleavage was detected between arginine and proline residues within the MUC1 variable number tandem repeat region (**fig. S3b**), which was consistent with previous reports that KLK5 preferentially cleaves substrates with arginine at the P1 position(*25*, *26*).

We next asked whether KLK5 could remodel mucins on intact cells. K562, Molm-13, and Nomo-1 cells treated with rhKLK5 had reduced the cell-surface mucin levels, although the reduction was smaller than that produced by StcE (**Fig. 2D**). Pretreatment with sialidase markedly reduced the KLK5-dependent loss of cell-surface mucin signal, indicating that the sialylation state of mucins influences KLK5-mediated cleavage. Increasing concentrations of rhKLK5 produced a corresponding decrease in mucin signal in K562 cells (**Fig. 2E and fig. S3C**).

The glycocalyx also contains large proteoglycans that can influence ligand binding to its intended receptor on the cell surface. For example, FGF2 binding to its receptor on the cell surface depends strongly on heparan sulfate proteoglycans(*27*, *28*). Therefore, we examined whether KLK5 affected heparan sulfate-dependent FGF2 binding. KLK5 reduced FGF2 binding in Capan-2, GSU, PDX1294, K562, Molm-13, and Nomo-1 cells (**Fig. 2F**). Across the six cell lines, KLK5 reduced FGF2 binding by approximately 56% relative to untreated cells, compared with a 98% reduction following heparinase treatment and an 11% reduction following chondroitinase treatment. In Capan-2, GSU and PDX1294 cells, heat inactivation of KLK5 attenuated this reduction (**fig. S3D**).

Finally, we measured whether KLK5 altered glycocalyx thickness using scanning angle interference microscopy (SAIM) (**Fig. 2G**). We previously used SAIM to measure glycocalyx thickness in MCF10A cells with doxycycline-titratable MUC1 expression(*10*). Using the same system, rhKLK5 reduced the average glycocalyx thickness by 14.7 nm in the absence of doxycycline and by 9.7 nm at 1 µg/ml doxycycline (**Fig. 2H and I**). rhKLK5 did not substantially alter the proliferation of GSU or NUGC4 cells at the concentrations used in subsequent experiments (**fig. S3E**). Overall, these results show that KLK5 remodels several protein components of the cancer-cell glycocalyx and reduces its nanoscale thickness.

## KLK5 increases MUC17 CAR-T cell avidity and antitumor activity

Having established that KLK5 remodels the cancer-cell glycocalyx, we next asked whether this activity could increase MUC17 CAR-T cell recognition of cancer cells (**Fig. 3A**). Using an acoustic force-based avidity assay, rhKLK5 treatment increased the fraction of MUC17 CAR-T cells that remained bound to ASPC-1 and NUGC4 cancer cells as the applied force was increased (**Fig. 3B and C**). At the plateau of the avidity curve, the fraction of MUC17 CAR-T cells bound to ASPC-1 cells increased from 22.5% to 30.6% following rhKLK5 treatment. StcE increased this binding to 48.2%, as expected due to its stronger mucin-cleaving activity. In NUGC4 cells, rhKLK5 increased the corresponding bound fraction from 28.9% to 35.0%. Therefore, rhKLK5 consistently increased MUC17 CAR-T cell avidity relative to untreated target cells, showing that KLK5-mediated glycocalyx remodeling was sufficient to strengthen CAR-T-cell binding avidity. Consistent with the increased avidity, combining rhKLK5 with MUC17 CAR-T cells increased GSU tumor-cell killing relative to MUC17 CAR-T cells alone (**Fig. 3D,E**). rhKLK5 alone did not substantially affect GSU cell growth, indicating that the increased killing was not due to a direct effect of KLK5 on tumor-cell proliferation.

**Fig. 3.**
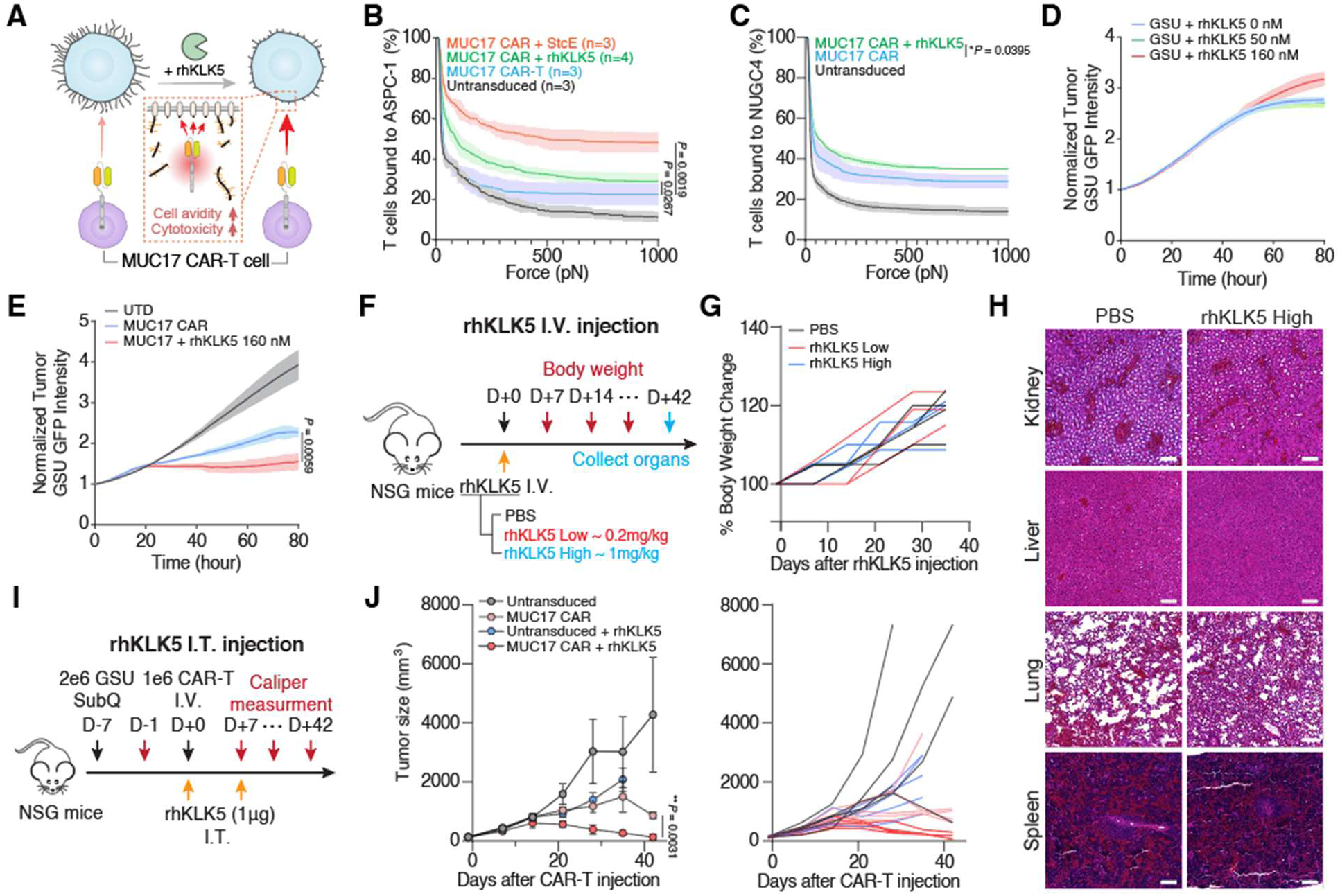
KLK5 enhances MUC17 CAR-T cell avidity and antitumor activity. (**A**) Schematic illustrating the effect of recombinant human KLK5 (rhKLK5) treatment on MUC17 CAR-T cell avidity and cytotoxicity. (**B** and **C**), Interaction strength between untreated, rhKLK5-treated or StcE-treated ASPC-1 (**B**) and NUGC4 (**C**) cells and either MUC17 CAR-T cells or untransduced T cells. The percentage of T cells remaining bound to target cells is shown as the applied acoustic force is increased from 0 to 1,000 pN. Results are mean ± s.d. from at least n = 3 independent measurements. (**D** and **E**) Representative real-time cytotoxicity assay of untransduced T cell (**D**) and MUC17 CAR-T cells (**E**) against untreated or rhKLK5-treated GSU cells at the indicated rhKLK5 concentrations and an 1:5 E:T ratio (relative to day 0 tumor seeding) from n = 3 three distinct human blood cell donors. Results are mean ± s.d. of n = 3 independent measurements. (**F**)-(**H**) Evaluation of systemic rhKLK5 administration *in vivo*. Three NOD.Cg-Prkdc^scid^ Il2rg^tm1Wjl^/SzJ (NSG) mice per group received the indicated intravenous (I.V.) injections of rhKLK5 or PBS (**F**), and body-weight change (**G**) and representative haematoxylin and eosin (H&E) staining of kidney, liver, lung and spleen (**H**) were assessed. Scale bar, 100 µm. (**I)** Experimental scheme for in vivo antitumor studies. Mice were subcutaneously injected with 2 × 10^6^ GSU cells on day -7. On day 0, mice received 1 × 10^6^ MUC17 CAR-T cells or untransduced T cells (n = 5 mice per group). Mice were further treated intratumorally (I.T.) with rhKLK5 (1 µg) on days 0 and 7. (**J**) Tumor growth curves (left) and individual mouse tumor growth trajectories (right). Results are mean ± s.e.m. of n = 5 mice per group.. In (**B**), (**E**), and (**J**), statistical analysis was performed by two-way ANOVA with correction for multiple comparisons.

We next examined the effects of rhKLK5 on CAR T cell function in vivo. To determine the tolerability of recombinant KLK5 protease, we first performed a safety assessment of systemically administered rhKLK5 alone in NSG mice (**Fig. 3F**). Because systemic dosing parameters for recombinant KLK5 have not been established, we evaluated a fivefold dose range of 0.2 and 1 mg/kg, with 1 mg/kg representing the highest practical dose using commercially available enzymatically active rhKLK5. The protease was well tolerated, as body weight remained stable over 42 days, and histological examination of the kidney, liver, lung and spleen did not reveal overt differences between PBS- and rhKLK5-treated mice (**Fig. 3G and H**). We subsequently tested rhKLK5 in a xenograft model of gastric cancer using the GSU cell line engrafted into the flank. Mice bearing established tumors received MUC17 CAR-T or untransduced T cells via IV injection on day 0, together with intratumoral injections of 1 µg rhKLK5 per mouse on days 0 and 7 (**Fig. 3I**). The combination of rhKLK5 and MUC17 CAR-T cells produced greater tumor control than MUC17 CAR-T cells alone (**Fig. 3J**). Thus, KLK5 enhanced MUC17 CAR-T cell activity both *in vitro* and *in vivo*. However, recombinant KLK5 required separate enzyme administration. We therefore asked whether KLK5 could instead be produced directly by the CAR-T cells.

## An ADAM10-based enzyme receptor enables membrane-tethered KLK5 delivery from CAR-T cells

We first investigated membrane-tethered KLK5 as a strategy to spatially restrict enzyme activity to the surface of the effector cell (**Fig. 4A**). Our initial design fused KLK5 to the CD28 transmembrane domain through a flexible Gly/Ser-rich extracellular linker. A shorter (GGGGS)_3_ linker did not support KLK5 expression, whereas extending the extracellular linker to 35 amino acids improved expression, indicating that KLK5 surface presentation was sensitive to the extracellular tether (**fig. S4A and B**).

**Fig. 4.**
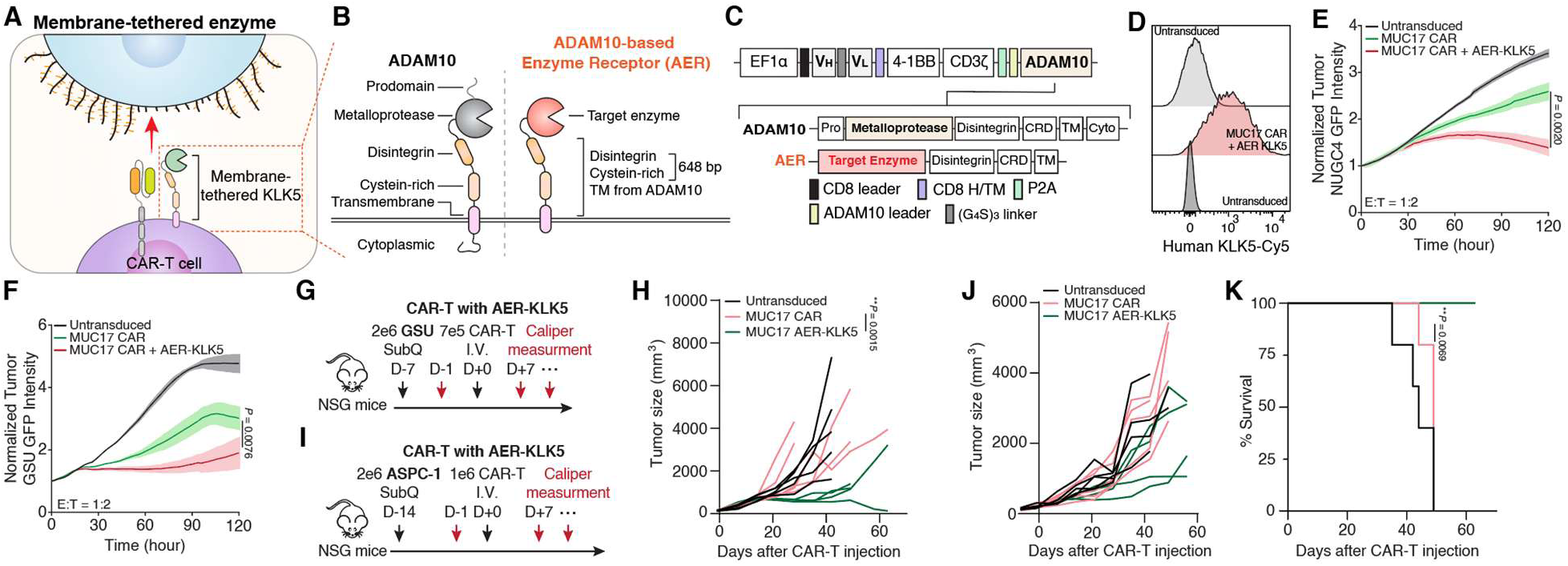
Membrane-tethered KLK5 enhances the antitumor activity of MUC17 CAR-T cells. **(A** and **B)**, Schematics illustrating the membrane-tethered enzyme strategy (**A**) and the endogenous ADAM10 structure together with the engineered ADAM10-based enzyme receptor (AER), generated by removing the prodomain and cytoplasmic domain of ADMA10 (**B**). (**C**) Graphic representation of the constructs used to generate CAR-T cells expressing AER-KLK5. (**D**) Flow cytometry analysis of human KLK5 expression on untransduced T cells or MUC17 CAR-T cells expressing AER-KLK5. (**E** and **F**) Representative real-time cytotoxicity assays of untransduced T cells, MUC17 CAR-T cells, and MUC17 CAR-T cells expressing AER-KLK5 against NUGC4 (**E**) and GSU (**F**) target cells at 1:2 E:T ratio (relative to day 0 tumor seeding) from n = 3 three distinct human blood cell donors. Results are mean ± s.d. of n = 3 independent measurements. (**G** and **H**) *in vivo* antitumor activity in the GSU xenograft model. NSG mice were subcutaneously injected with 2 × 10^6^ GSU cells on day -7. On day 0, mice received 0.7 × 10^6^ MUC17 CAR-T cells or untransduced T cells (n = 5 mice per group) (**G**), and tumor growth was monitored over time (**H**). (**I**)**-**(**K**) *in vivo* antitumor activity in the ASPC-1 xenograft model. NSG mice were subcutaneously injected with 2 × 10^6^ ASPC-1 cells on day -14. On day 0, mice received 1 × 10^6^ MUC17 CAR-T cells or untransduced T cells (n = 5 mice per group) (**I**). Tumor growth curves (**J**) and survival percentage (**K**) are shown. In (**E**), (**F**), and (**H**), statistical analysis was performed by two-way ANOVA with correction for multiple comparisons. In (**K**), statistical analysis was performed using a log-rank (Mantel-Cox) test.

To improve membrane presentation, we considered the architecture of endogenous membrane proteases. ADAM10 is a membrane-anchored metalloprotease with an extended extracellular structure containing disintegrin and cysteine-rich regions between its catalytic module and transmembrane segment(*29*). Structural studies further demonstrate that the membrane-associated organization of ADAM10 controls the positioning of proteolytic activity relative to membrane-proximal substrates(*30*). Therefore, we repurposed the membrane-tethering architecture of ADAM10 as a scaffold for enzyme presentation. We removed the ADAM10 prodomain and cytoplasmic domain and replaced its native catalytic module with KLK5, while retaining the extended ADAM10-derived extracellular and transmembrane regions, to generate an ADAM10-based enzyme receptor that we termed AER-KLK5 (**Fig. 4B and C**). The retained ADAM10-derived extracellular region comprises approximately 216 amino acids, providing substantially greater spacing than the 35-amino-acid linker used in the CD28-based construct.

AER-KLK5 exhibited higher expression than the CD28-based membrane-tethered KLK5 construct in Jurkat cells (**fig. S4B**). To determine whether the AER architecture could support other proteases, we generated constructs containing ADAM10, cathepsin K, human neutrophil elastase, or KLK5. ADAM10 mediates selective shedding of transmembrane mucins and mucin-like glycoproteins(*31*), cathepsin K degrades cell-surface mucins and proteoglycans(*23*), and human neutrophil elastase has been reported to proteolytically release cell-surface mucins, including MUC16(*32*). Therefore, we compared AER constructs incorporating these proteases with AER-KLK5. Among the constructs tested, AER-KLK5 produced the highest surface expression in Jurkat cells (**fig. S4C and D**). In primary human T cells, AER-KLK5 again produced the highest detectable enzyme expression among the tested AER constructs when combined with the MUC17 CAR (**fig. S4E**). Following coculture with NUGC4 cells for 24 h, AER-KLK5 MUC17 CAR-T cells produced a modest reduction in the cell-surface mucin signal on target cells (**fig. S4G and H**). Surface staining with an anti-KLK5 antibody confirmed detectable KLK5 on MUC17 CAR-T cells expressing AER-KLK5 (**Fig. 4D**). We then tested whether membrane-tethered KLK5 improved CAR-T cell cytotoxicity. In real-time cytotoxicity assays, AER-KLK5 MUC17 CAR-T cells showed greater killing of both NUGC4 and GSU cells than conventional MUC17 CAR-T cells. AER-KLK5 also produced greater cytotoxicity than the CD28-based KLK5 display construct under matched conditions (**Fig. 4E, F, and** **fig. S4G and H**).

The activity of AER-KLK5 was subsequently evaluated *in vivo*. In the GSU xenograft model, mice bearing established tumors received 0.7 × 10^6^ untransduced T cells, conventional MUC17 CAR-T cells, or AER-KLK5 MUC17 CAR-T cells (**Fig. 4G**). MUC17 CAR-T cells with AER-KLK5 produced greater tumor control than conventional MUC17 CAR-T cells (**Fig. 4H**). We subsequently tested the strategy in the ASPC-1 xenograft model using 1 × 10^6^ CAR-T cells (**Fig. 4I**). Tumor responses were more heterogeneous in this model, with pronounced tumor control observed in a subset of AER-KLK5-treated animals (**Fig. 4J**). Nevertheless, AER-KLK5 MUC17 CAR-T cells prolonged survival relative to conventional MUC17 CAR-T cells (**Fig. 4K**). These results show that membrane-tethered KLK5 can improve MUC17 CAR-T cell activity *in vivo*, including prolonged survival in the ASPC-1 model.

## KLK5-secreting CAR-T cells enhance antitumor activity across multiple tumor models

The context-dependent activity of membrane-tethered KLK5 suggested that restricting the enzyme to the T-cell surface may limit its access to glycocalyx substrates beyond the area of direct tumor-cell contact. Therefore, we developed a complementary strategy in which CAR-T cells secrete KLK5 into the extracellular environment (**Fig. 5A**). Using a P2A-based bicistronic construct containing an Igκ secretion leader, we generated MUC17 CAR-T cells that co-express secreted KLK5, referred here to as MUC17 kCAR-T cells (**Fig. 5B**).

**Fig. 5.**
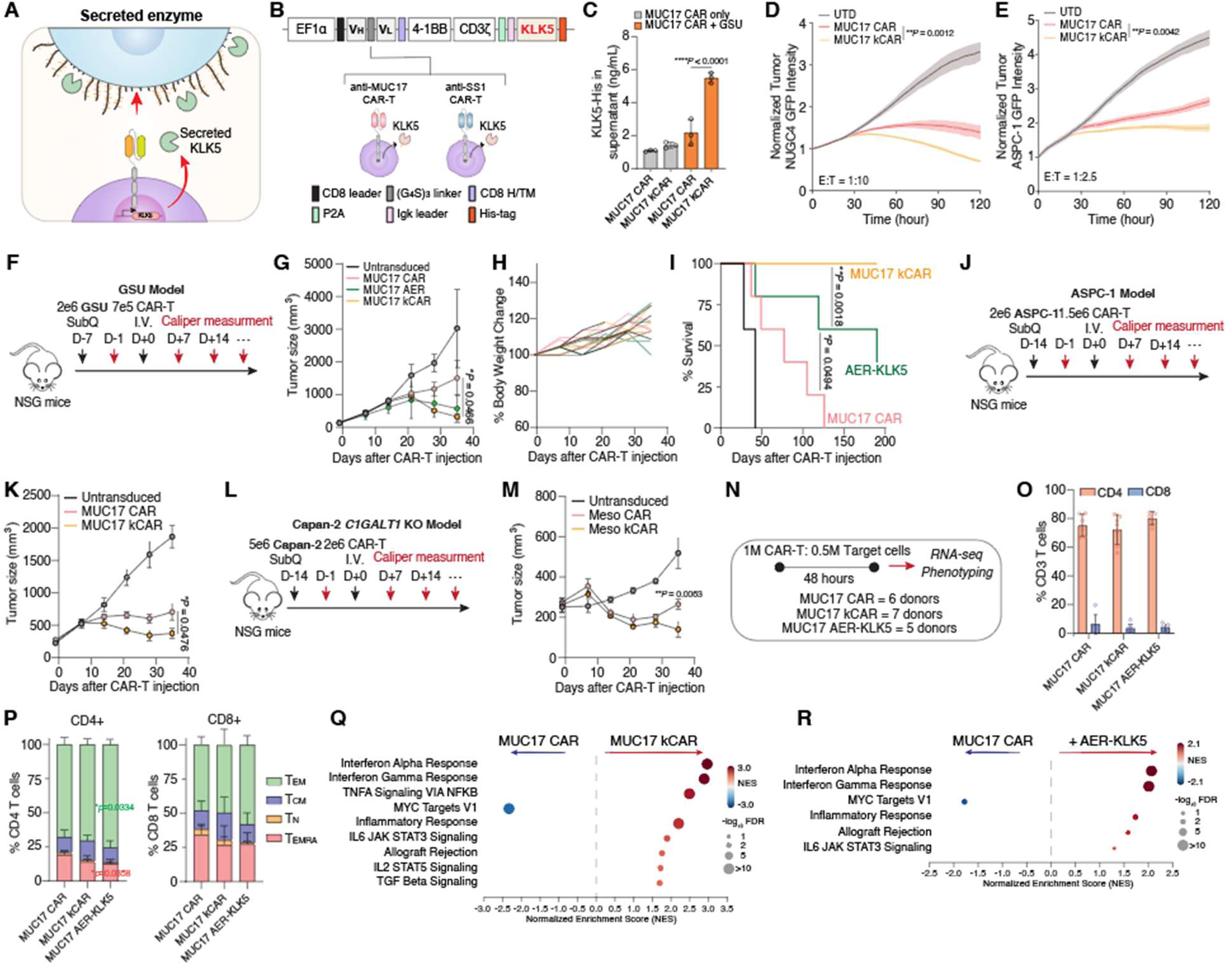
CAR-T cell secretion of KLK5 enhances antitumor activity and promotes an activated transcriptional state. **(A)** Schematic illustrating the secreted-enzyme strategy for local remodeling of the tumor-cell glycocalyx by CAR-T cells. (**B)** Graphic representation of the constructs used to generate MUC17- or mesothelin-targeting CAR-T cells secreting KLK5 (kCAR). (**C)** Quantification of secreted KLK5 by His-tag ELISA in supernatants from MUC17 CAR-T and MUC17 kCAR-T cells cultured alone or following coculture with GSU cells at 2:1 E:T ratio. Results are mean ± s.d. of n = 3 distinct human blood-cell donors. (**D** and **E)** Representative real-time cytotoxicity assays of untransduced T cells, MUC17 CAR-T cells and MUC17 kCAR-T cells against NUGC4 (**D**) and ASPC-1 (**E**) cells at E:T ratios of 1:10 and 1:25, respectively (relative to day 0 tumor seeding). Data are representative of experiments performed with n = 3 distinct human blood-cell donors, with results shown as mean ± s.d. of n =3 independent measurements. (**F**)-(**I)** *in vivo* antitumor activity in the GSU xenograft model. NSG mice were subcutaneously injected with 2 × 10^6^ GSU cells on day -7. On day 0, mice received 0.7 × 10^6^ untransduced T cells, MUC17 CAR-T cells, MUC17 kCAR-T cells or MUC17 CAR-T cells expressing AER-KLK5 (n = 5 mice per group) (**F**). Tumor growth (**G**), body-weight change (**H**) and survival percentage (**I**) were monitored over time. Tumor-growth data are mean ± s.e.m. of n = 5 mice per group. (**J** and **K**) *in vivo* antitumor activity in the ASPC-1 xenograft model. NSG mice were subcutaneously injected with 2 × 10^6^ ASPC-1 cells on day -14. On day 0, mice received 1.5 × 10^6^ untransduced T cells, MUC17 CAR-T cells or MUC17 kCAR-T cells (n = 5 mice per group) (**J**), and tumor growth was monitored over time (**K**). Results are mean ± s.e.m. of n = 5 mice per group. (**L** and **M**) *in vivo* antitumor activity in the Capan-2 *C1GALT1*-knockout xenograft model. NSG mice were subcutaneously injected with 5 × 10^6^ Capan-2 *C1GALT1* KO cells on day -14. On day 0, mice received 2 × 10^6^ untransduced T cells, mesothelin-targeting CAR-T cells or mesothelin-targeting kCAR-T cells (n = 5 mice per group) (**L**), and tumor growth was monitored over time (**M**). Results are mean ± s.e.m. of n = 5 mice per group. (**N**) Schematic of the experimental workflow for T-cell phenotyping and RNA-sequencing analysis. The indicated T-cell populations were cocultured with GSU cells at an E:T ratio of 2:1 for 48 h before analysis. Experiments included MUC17 CAR-T cells, MUC17 kCAR-T cells and MUC17 CAR-T cells expressing AER-KLK5 from n = 6, 7 and 5 distinct human blood-cell donors, respectively. (**O** and **P**) CD4 and CD8 population (**O**) and phenotype (**P**) of indicated CAR-T cells. Cells were grouped by flow cytometry according to T-cell phenotypes as follows: naïve (T_N_): CCR7^+^CD45RO^-^, T_CM_: CCR7^+^CD45RO^+^, T_EM_: CCR7^-^CD45RO^+^ and effector (T_E_): CCR7^-^CD45RO^-^. Results are mean ± s.d. of n = 6, 7 and 5 distinct human blood-cell donors, respectively. (**Q** and **R**) Gene-set enrichment analysis of RNA-sequencing data comparing MUC17 kCAR-T cells with MUC17 CAR-T cells (**Q**) and MUC17 CAR-T cells expressing AER-KLK5 with MUC17 CAR-T cells (**R**) after 48 h of coculture with GSU cells. Selected Hallmark gene sets are shown. The horizontal axis indicates the normalized enrichment score (NES), with positive values indicating enrichment in KLK5-engineered CAR-T cells and negative values indicating enrichment in conventional MUC17 CAR-T cells. Dot color indicates NES and dot size represents -log_10_(FDR). In (**C**), statistical analysis was performed by one-way ANOVA with Tukey’s post hoc tests. In (**D**), (**E**), (**G**), (**K**), and (**M**), statistical analysis was performed by two-way ANOVA with correction for multiple comparisons. In (**I**), statistical analysis was performed using a log-rank (Mantel-Cox) test.

We first tested whether KLK5 was released from MUC17 kCAR-T cells. A His tag was incorporated into KLK5, allowing extracellular KLK5 to be quantified by ELISA. His-tagged KLK5 was readily detected in supernatants from MUC17 kCAR-T cells. Following coculture with GSU target cells, extracellular KLK5 increased approximately 3.9-fold, from 1.42 to 5.50 ng/ml, compared with kCAR-T cells cultured without tumor cells (**Fig. 5C**). We next compared conventional MUC17 CAR-T and MUC17 kCAR-T cells. Across independent human blood-cell donors, MUC17 kCAR-T cells showed substantially greater tumor-cell killing than conventional MUC17 CAR-T cells against both NUGC4 and ASPC-1 cells in real-time cytotoxicity assays (**Fig. 5D and E**). These data indicated that secreted KLK5 enhanced MUC17 CAR-T cell activity across gastric and pancreatic tumor cell contexts.

The membrane-tethered and secreted KLK5 strategies were then compared directly *in vivo*. In the GSU xenograft model, mice received untransduced T cells, conventional MUC17 CAR-T cells, AER-KLK5 MUC17 CAR-T cells, or MUC17 kCAR-T cells (**Fig. 5F**). MUC17 kCAR-T cells produced the strongest tumor control among the tested CAR-T cell groups and significantly improved tumor control relative to conventional MUC17 CAR-T cells (**Fig. 5G**). Body weight remained stable across treatment groups (**Fig. 5H**), and mice treated with MUC17 kCAR-T cells exhibited markedly prolonged survival compared to conventional MUC17 CAR-T and AER-KLK5 CAR-T cells (**Fig. 5I**).

We observed a similar improvement in the ASPC-1 pancreatic cancer xenograft model. Mice bearing established ASPC-1 tumors received conventional MUC17 CAR-T cells, MUC17 kCAR-T cells, or untransduced T cells (**Fig. 5J**). As observed in the GSU model, MUC17 kCAR-T cells produced greater tumor control than conventional MUC17 CAR-T cells (**Fig. 5K**). To determine if this approach modified MUC17 antigen expression in the tumor, we analyzed the residual tumors. MUC17 staining was reduced following MUC17 CAR-T cell treatment relative to tumors treated with untransduced T cells (**fig. S5A and B**). Since CAR-T cell exposure itself reduced MUC17 staining, residual MUC17 levels could not be interpreted as a specific readout of KLK5 activity *in vivo*.

To determine whether the effect of KLK5 extended beyond MUC17 targeting, we generated mesothelin CAR-T cells secreting KLK5 and compared their activity with conventional mesothelin CAR-T cells against Capan-2 pancreatic cancer cells with *C1GALT1* KO. We previously established this model to increase truncated O-glycan and Tn-MUC1 expression on Capan-2 cells, providing a glycan-rich tumor model for evaluating CAR-T cell activity(*22*). Mesothelin kCAR-T cells showed greater real-time cytotoxicity than conventional mesothelin CAR-T cells (**fig. S5D**). In the Capan-2 *C1GALT1* KO xenograft model, mesothelin kCAR-T cells similarly produced greater tumor control than conventional mesothelin CAR-T cells (**Fig. 5L and M**). Thus, the effect of KLK5 was not restricted to MUC17-directed CAR-T cells.

We also tested KLK5 secretion with a Tn-MUC1-targeting CAR. In contrast to the membrane-proximal MUC17 CAR, the Tn-MUC1 CAR recognizes glycopeptide epitopes contained within the protease-accessible MUC1 tandem-repeat region(*13*). KLK5 secretion produced only a modest improvement in Tn-MUC1 CAR-T cell cytotoxicity *in vitro* and did not produce a clear improvement *in vivo* (**fig. S5E-G**). Due to the fact that KLK5 directly cleaves the MUC1 VNTR, proteolytic remodeling in this setting may simultaneously reduce the physical glycocalyx barrier and remove CAR-binding sites. This result was consistent with the epitope-position-dependent model established in **Fig. 1**. To identify KLK5 variants with greater activity, we screened AI-guided KLK5 variants, but none produced a clear improvement in CAR-T cell killing compared with wild-type KLK5 (**fig. S6**).

We next examined whether KLK5 engineering altered the state of CAR-T cells following tumor-cell engagement. MUC17 CAR-T, MUC17 kCAR-T, and AER-KLK5 MUC17 CAR-T cells from n = 6, 7, and 5 distinct human blood-cell donors were cocultured with GSU cells for 48 hours before flow-cytometric phenotyping and RNA-sequencing analysis (**Fig. 5N**). We did not observe substantial differences in the overall CD4 ratio or most of the measured T-cell differentiation states by flow cytometry (**Fig. 5O and P**). AER-KLK5 CAR-T cells showed a modest increase in effector-memory cells and a corresponding decrease in terminally differentiated effector-memory cells.

RNA-sequencing analysis of CAR-T cells isolated after tumor-cell coculture showed broader transcriptional differences. KLK5-engineered CAR-T cell groups were enriched for interferon-α response, interferon-γ response, inflammatory response, allograft rejection, and IL6-JAK-STAT3 signaling relative to conventional MUC17 CAR-T cells, regardless of how the enzyme was expressed (**Fig. 5Q and R**). MUC17 kCAR-T cells showed additional enrichment of TNFα signaling through NF-κB and IL2-STAT5 signaling, whereas conventional MUC17 CAR-T cells were relatively enriched in MYC Targets V1. These data indicate that KLK5-engineered CAR-T cells acquired distinct, activation-associated transcriptional states during tumor-cell coculture. Because unstimulated KLK5-expressing CAR-T cells were not profiled, we cannot distinguish direct transcriptional effects of KLK5 expression from changes arising secondarily from differences in tumor-cell engagement or cytotoxic activity. Nevertheless, across the functional studies, secreted KLK5 produced the most consistent enhancement of CAR-T cell antitumor activity, supporting local glycocalyx remodeling as a strategy to improve CAR-T cell function.

## Discussion

In summary, we show that the mucin-rich glycocalyx regulates CAR-T cell activity through opposing effects on antigen availability and physical resistance to cell-cell engagement. Increasing mucin expression initially enhanced the activity of a mucin-targeting CAR, whereas further increases in mucin density reduced cytotoxicity despite greater target expression. Proteolytic remodeling of the mucin layer increased recognition of membrane-proximal MUC1 and MUC17 epitopes and enhanced CAR-T cell avidity and killing. We further identified KLK5 as a human protease that remodels the cancer-cell glycocalyx and engineered CAR-T cells to either display or secrete KLK5. Both approaches enhanced CAR-T cell activity, with secreted KLK5 producing the strongest tumor control and survival benefit among the CAR-T cell formats tested *in vivo*. Together, these results extend our previous studies of the glycocalyx as a physical barrier to immune-cell attack and show that engineered T cells can directly modify this barrier(*10*).

Our results indicate that cell-surface mucin expression can have two opposing effects on mucin-targeting CAR-T cells. Increasing expression of a mucin antigen increases the number of available CAR-binding sites and can enhance CAR-T cell activity. However, increasing mucin density also expands and crowds the glycocalyx, which can resist close membrane apposition. Thus, high expression of a mucin target does not necessarily result in greater CAR-T cell activity. This effect is also not restricted to CARs that directly recognize mucins. In our previous studies, increasing mucin expression reduced the activity of CARs directed against other membrane proteins(*10*). Disruption of tumor-cell *N*-glycans has similarly been shown to improve immune-synapse formation and CAR-T cell cytotoxicity(*12*). More recently, CD43 was identified as a large glycoprotein barrier that can protect leukemia cells from immune-cell attack(*24*). These observations suggest that mucins and other bulky glycans and glycoproteins in the glycocalyx can interfere with CAR-T cell recognition even when they are not the CAR target.

We also found that the effect of glycocalyx remodeling on CAR-T cell function depended on the position of the targeted epitope. StcE treatment reduced recognition of epitopes within the cleaved MUC1 tandem-repeat region but increased accessibility to membrane-proximal MUC1 and MUC17 epitopes. Previous studies have also demonstrated that CAR activity can depend on the position of the targeted epitope, with membrane-proximal epitopes favoring close membrane apposition and exclusion of CD45 from the receptor-binding interface(*14*). Consistent with this idea, KLK5 improved MUC17- and mesothelin-directed CAR-T cell activity, whereas the improvement was limited for a Tn-MUC1 CAR, whose target lies within the KLK5-sensitive MUC1 tandem-repeat region. Thus, removing a mucin barrier may be beneficial when the CAR target is preserved, but less effective when the enzyme also removes the targeted epitope. Therefore, the effect of glycocalyx remodeling depends on both the amount of mucin present and whether the targeted epitope remains accessible after proteolysis.

We identified KLK5 activity against MUC1 through a focused screen of kallikrein-related proteases. However, KLK5 is unlikely to be the only human protease that can be used for this purpose. Many extracellular proteases remain poorly tested for their ability to remodel the cancer-cell glycocalyx. Recent work identified cathepsin K as a human protease capable of degrading several components of the glycocalyx and reducing glycocalyx thickness(*23*). Our results show that KLK5 can cleave several large glycocalyx proteins, including MUC1, MUC16, MUC17 and CD43. In contrast, PSGL-1 and CD45 were less affected under the same conditions, indicating that KLK5 does not cleave all highly glycosylated surface proteins equally. KLK5 also reduced FGF2 binding and decreased glycocalyx thickness measured by SAIM, suggesting that its effects extend beyond a single mucin substrate. A broader comparison of human extracellular proteases may identify enzymes with different substrate preferences or greater activity against specific tumor glycocalyces.

Secreted KLK5 improved CAR-T cell activity more consistently than membrane-tethered KLK5 and produced the strongest *in vivo* effects in the models tested. More extracellular KLK5 was detected after coculture with antigen-positive tumor cells, although the construct itself was not antigen regulated. One possible explanation for the greater activity of secreted KLK5 is broader access to tumor cell glycocalyces beyond sites of direct T cell contact. We did not directly measure KLK5 distribution within tumors, so this remains a possible explanation rather than an established mechanism. Systemically administered rhKLK5 did not produce detectable weight loss or histopathological changes at the doses tested, supporting further evaluation of secreted KLK5 delivery. If tighter control is required, KLK5 production could be linked to tumor-antigen recognition using synthetic receptor circuits (*33–35*) Alternatively, KLK5 could be targeted directly to tumor cells, as demonstrated for antibody-targeted sialidase and StcE constructs(*16*, *17*).

The contribution of individual KLK5 substrates to improved CAR-T cell activity remains to be defined. Because KLK5 acts on several glycocalyx proteins, the most useful protease will likely depend on both the composition of the tumor glycocalyx and the position of the CAR target epitope. Overall, our results show that CAR-T cells can be engineered not only to recognize tumor antigens but also to modify the glycocalyx encountered during tumor-cell engagement. Therefore, this engineering approach provides a direct strategy to improve CAR-T cell access and activity in mucin-rich solid tumors.

## Supporting information

Park Supplementary Information

## Acknowledgments

We thank the core facilities at the MGH Cancer Center: Flow Cytometry, Histopathology, and Blood Bank. We thank Lauren Pepi and Richard Cummings at the Beth Israel Deaconess Medical Center Glycomics Center for assistance with glycomics analysis. This work was supported by NIH R01 CA238268 (M.V.M.) and NIH K99CA303346 (S.P.). S.P. was also supported by the Damon Runyon Cancer Research Foundation as a Merck Fellow (DRG-2529-24).

## Author Contributions

S.P. designed the research and performed experiments. All authors made a significant contribution to the discussion. S.P., T.R.B., and M.V.M. wrote the manuscript with feedback from all authors.

## Data, code, and materials availability

All data shown in this manuscript are provided as source data files. All additional data and materials that can be shared will be released using a material transfer agreement. Requests should be directed to Sangwoo Park or Marcela V. Maus.

## Supplementary Materials

Materials and Methods, Supplementary Note 1, Figs. S1 to S6, Tables S1

