## Supplementary material for "Engineering CAR-T cells to remodel the mucin-rich cancer cell glycocalyx": Park Supplementary Information

**This Supplementary Information file contains:**

**- Materials and Methods**

**- Supplementary Note 1**

**- Supplementary Figures 1-6**

**- Supplementary Table 1**

### Materials and Methods

#### Primary cells, cell lines, PDX tumor, engineered cell lines

Capan-2 (ATCC; HTB-80), GSU, NUGC4, ASPC-1, K562, Molm-13, Nomo-1 and Jurkat (ATCC; TIB-152) cells were cultured in RPMI 1640 media (Thermo Fisher Scientific; Cat# 72400047) supplemented with 10% fetal bovine serum (Thermo Fisher Scientific) and 1x penicillin/streptomycin (Thermo Fisher Scientific) at 37°C in 5% $CO_2$. MCF10A cells were cultured in Dulbecco's modified Eagle's medium (DMEM)/F12 medium (Thermo Fisher Scientific) supplemented with 5% horse serum (Thermo Fisher Scientific), 20 ng/ml epidermal growth factor (PeproTech), 10 mg/ml insulin (Sigma), 500 ng/ml hydrocortisone (Sigma), 100 ng/ml cholera toxin (Sigma) and 1x penicillin/streptomycin (Thermo Fisher Scientific)) at 37°C in 5% $CO_2$. The non-immortalized pancreatic ductal adenocarcinoma PDX cell line PDX1294 was kindly provided by the Liss laboratory at Massachusetts General Hospital and cultured in DMEM/F12 medium supplemented with 10% FBS and 1x penicillin/streptomycin. Cell lines were transduced to express click beetle green (CBG) luciferase and enhanced GFP (eGFP), then sorted using a BD FACSAria to isolate a clonal or different population of transduced cells.

Human T cells were isolated (Stem Cell Technologies, 15061) from healthy donor leukopaks obtained through the Massachusetts General Hospital Blood Bank using the StemCell Technologies T-cell RosetteSep Isolation Kit (Cat. no. 15061). T cells were cultured in RPMI 1640 media supplemented with 10% fetal bovine serum, 20 U/ml recombinant human IL-2 (PeproTech), and 1x penicillin/streptomycin at 37°C in 5% $CO_2$. Donor recruitment, screening and written informed consent were conducted by the MGH Blood Bank. Investigators received only de-identified samples and had no access to donor information.

#### Doxycycline-inducible MUC1 cell model

The doxycycline-inducible MUC1-GFP MCF10A model was generated as previously described(*7*, *10*). Briefly, MUC1-GFP containing 42 tandem repeats was stably expressed under control of a tetracycline-inducible promoter, and the clonally expanded 1E7 line was selected for its titratable MUC1 surface expression. The *C1GALT1* KO derivative used in this study was generated previously from the 1E7 clone by CRISPR/Cas9-mediated disruption of *C1GALT1*, followed by clonal expansion. The resulting clone, 2A7, retains doxycycline-titratable MUC1 expression in a C1GALT1-deficient background. This cell line was generated and characterized in our previous study(*10*). For cell-surface MUC1 density experiments, cells were treated with the indicated concentrations of doxycycline for 24 h before subsequent enzyme treatment, flow cytometric analysis or cytotoxicity assays.

**In vivo models**

Male and female NOD.Cg-Prkdcscid Il2rgtm1Wjl/SzJ (NSG) mice, 6-11 week-old, were bred in-house were bred in-house and maintained under specific pathogen-free conditions at the MGH Center for Cancer Research. Mice were housed at temperatures of 21.1-24.5°C (70-76°F), 30-70% humidity, and a 12:12 light-dark cycle. All animal studies were performed in accordance with protocols approved by the Massachusetts General Hospital Institutional Animal Care and Use Committee (IACUC; 2020N000114). The maximal tumor size permitted (20 mm in diameter for a single tumor or 10 mm for two tumors) was not exceeded in any experiment. All animal procedures, including injections and monitoring, were carried out by three animal technicians who were independent of the study hypotheses and blinded to treatment groups. Both male and female mice were used, and animals were randomized before treatment to obtain comparable tumor sizes across experimental groups. For subcutaneous tumor models, the indicated tumor cells were suspended in 100 µL of a 1:1 mixture of PBS and Matrigel (Corning) and injected into each mouse subcutaneously. CAR-T cells or donor-matched untransduced T cells were administered intravenously through the tail vein at the indicated time points and doses in 100 µL PBS. Caliper measurements were taken weekly. For the IHC study, mice were euthanized at the indicated time points to assess T cell infiltration by immunohistochemistry. Mice were monitored until the experimental endpoint or until euthanasia criteria defined by the approved IACUC protocol were reached. Tumor volume was calculated using the standard formula: Volume = (length × width$^2$) / 2. For GSU xenograft studies, 2 x $10^6$ tumor cells were implanted subcutaneously 7 days before T-cell administration. For ASPC-1 studies, 2 x $10^6$ cells were implanted 14 days before treatment, and for Capan-2 *C1GALT1* KO studies, 5 x $10^6$ cells were implanted 14 days before treatment. The number of CAR-T or untransduced T cells administered varied between experiments and is indicated in the corresponding figure legends. In experiments using recombinant KLK5, mice received 1 µg rhKLK5 per mouse by intratumoral injection on days 0 and 7.

**Generation of CAR constructs**

MUC17-, mesothelin- and Tn-MUC1-targeting CAR constructs were synthesized and cloned into a third-generation lentiviral backbone under the control of the human EF-1α promoter (GenScript). The mesothelin- and Tn-MUC1-targeting scFvs were derived from SS1 and 5E5, respectively, whereas the MUC17 CAR was generated as previously described(*13*, *18*, *38*). All CAR constructs contained a CD8 hinge and transmembrane domain, a 4-1BB co-stimulatory domain and an intracellular CD3ζ signaling domain.

To generate membrane-tethered KLK5, human KLK5 was first fused to a CD28 transmembrane domain using Gly/Ser-rich extracellular linkers of different lengths. We subsequently developed an ADAM10-based enzyme receptor (AER) to improve surface presentation of KLK5. For AER-KLK5, the prodomain and cytoplasmic domain of ADAM10 were removed and its metalloprotease domain was replaced with human KLK5, while retaining the ADAM10-derived extracellular scaffold and transmembrane region. AER constructs containing KLK5, ADAM10, cathepsin K or human neutrophil elastase were generated using the same architecture. For secretion of KLK5, an Igκ leader peptide was placed upstream of human KLK5 and the resulting secretion cassette was linked to the CAR through a P2A sequence. These constructs are referred to as kCARs. MUC17, mesothelin and Tn-MUC1 kCAR constructs were generated using the corresponding CAR backbones. A His tag was incorporated into the secreted KLK5 construct to allow detection of extracellular KLK5. To generate antigen-targeted KLK5, secreted KLK5 was fused to scFvs recognizing CD19, mesothelin or Tn-MUC1. The CD19-, mesothelin- and Tn-MUC1-targeting scFvs were derived from blinatumomab, SS1 and 5E5, respectively. All constructs were verified by sequencing before lentiviral production.

**CAR T cell production**

Leukapheresis product from anonymous healthy human donors was purchased from the MGH blood bank, under an institutional review board-exempt protocol. Donor recruitment, screening, and written informed consent were conducted by the MGH Blood Bank. The investigators received only de-identified samples and had no access to donor information. Stem Cell Technologies T cell Rosette Sep Isolation kit was used to isolate T cells. Bulk human T cells were activated on Day 0 using CD3/CD28 Dynabeads (Life Technologies) at a 1:3 T cell:bead ratio to generate CAR T cells and untransduced T cells from the same donors to serve as controls. T cells were grown in RPMI 1640 media with GlutaMAX and HEPES supplemented with 10% FBS, penicillin, streptomycin, and recombinant human IL-2 (20 IU per ml;

Peprotech). On Day 1 (24 h after activation), cells were transduced with CAR lentivirus at an MOI of 5-10. CAR-T cells were expanded with IL-2 containing cell culture media addition every 2-3 days to main the concentration between 0.5-1 x $10^6$ cells/mL. Dynabeads were removed via magnetic separation on Day 6, and cells were assessed by flow cytometry with mCherry expression, ALFA-tag, or anti-$(G_4S)_3$ linker antibody binding on Days 12-14 to determine transduction efficiency prior to cryopreservation. Prior to use in *in vitro* and *in vivo* functional assays, CAR T cells and untransduced cells were thawed and rested for 18-24 h in the presence of IL-2.

**Scanning angle interference microscopy**

Scanning angle interference microscopy (SAIM) was performed using the same Ring-SAIM system and analysis pipeline described previously(*7*, *10*, *39*). Briefly, Bzip-expressing MCF10A cells expressing doxycycline-inducible MUC1-GFP were seeded on fibronectin-functionalized silicon chips at 2 x $10^4$ cells $cm^{-2}$ and cultured with the indicated doxycycline concentrations for 24 h. Cells were treated with rhKLK5 or vehicle under the indicated conditions before imaging. Cell-surface was labeled with Alexa Fluor 647-Azip, a cognate leucine zipper, and live cells were imaged at 37 °C. Glycocalyx thickness was determined from the reconstructed height of the AF647-Azip signal relative to the fluorescent reference layer on the silicon substrate, as previously described.

**Recombinant enzyme treatment and glycocalyx analysis**

Recombinant human KLK5 (rhKLK5; R&D Systems, 1108-SE-010) was used for cell-surface glycocalyx-remodeling experiments. Unless otherwise indicated, rhKLK5 was diluted 1:10 or 1:20 from the supplied protein preparation into the corresponding cell-culture medium and incubated with cells at 37 °C for 1-2 hours. For cytotoxicity experiments, rhKLK5 was added at the initiation of CAR-T cell and tumor-cell coculture and remained present throughout the assay. StcE mucinase (Sigma-Aldrich, SAE0202) was used at 100 nM and incubated with cells for 1 h at 37 °C before washing and downstream analysis. For desialylation experiments, neuraminidase from vibrio cholerae (Sigma-Aldrich, 11080725001) was diluted 1:10 and incubated with cells at 37 °C for 2 h before subsequent rhKLK5 treatment. To compare the mucin-cleaving activity of kallikrein-related proteases, recombinant human KLK4, KLK5, KLK6, KLK7 and KLK13 (R&D Systems) were incubated with the indicated substrates under matched conditions.

Proteolysis reactions were performed in 0.1 M $NaH_2PO_4$, pH 7.4, for 16 h at 37 °C. KLKs were used at a final concentration of 1.6 μM and an enzyme-to-substrate molar ratio of 1:50. Reaction products were analyzed by SDS-PAGE and western blotting. For purified mucin cleavage assays, recombinant human MUC17 (ACROBiosystems, MU7-H52H3) or MUC16 (ACROBiosystems, CA5-H82F8) was incubated with rhKLK5 under the same reaction conditions and analyzed by Coomassie staining or western blotting as indicated. To assess whether KLK5-dependent changes in the glycocalyx required proteolytic activity, rhKLK5 was heat-inactivated at 90 °C for 30 min. Active and heat-inactivated rhKLK5 were diluted identically and applied to cells under otherwise matched conditions. Changes in heparan sulfate-dependent ligand binding were evaluated by FGF2 staining following enzymatic treatment. Capan-2, GSU, PDX1294, K562, Molm-13 and Nomo-1 cells were treated with rhKLK5, heparinase I or chondroitinase ABS at 37 °C for 2 h. Heparinase I (New England Biolabs, P0735S) was prepared as a 1,200 U/ml 10x working solution and used at a final concentration of 120 U/ml. Chondroitinase ABS (Sigma-Aldrich, C3667-5UN) was used at a final concentration of 1 U/ml. For Capan-2, GSU and PDX1294 cells, active rhKLK5 was directly compared with an equivalent concentration of heat-inactivated rhKLK5. Following enzyme treatment, cells were washed and incubated with His-tagged recombinant human FGF2 (ACROBiosystems, FGC-H81E3-25ug) at a 1:100 dilution in FACS buffer. Cell-associated FGF2 was detected using an anti-His-tag antibody at a 1:200 dilution and quantified by flow cytometry. Chondroitin sulfate was measured using an anti-chondroitin sulfate antibody (Thermo Fisher Scientific, MA1-83055) diluted 1:100, followed by the appropriate anti-mouse secondary antibody. Flow-cytometric signals were normalized to the corresponding untreated controls where indicated.

**Flow cytometric analysis**

Flow cytometry was used to assess CAR-T cell phenotype, CAR expression and cell-surface glycocalyx components. The following antibodies were used for CAR-T cell analysis: Alexa Fluor 647 conjugated $(G4S)_3$ (E7O2V) antibody (69782S; Cell Signaling Technology; 1:100), Alexa Fluor 700 conjugated anti-human CD3 antibody (300424; BioLegend; 1:100), Per/Cy7 conjugated anti-human CD4 antibody (300518; BioLegend; 1:100), PerCP conjugated anti-human CD8α antibody (301032; BioLegend; 1:100), FITC conjugated anti-human CCR7 antibody (561271; BD BioSciences; 1:50), Brilliant Violet 421 conjugated anti-human CD45RA antibody (304130; BioLegend; 1:100), Alexa Fluor 700 conjugated anti-human CD45 antibody (304024; BioLegend; 1:100), and APC conjugated anti-human CD69 (310910;

BioLegend). Each antibody was diluted in PBS containing 2% FBS, as indicated. Adherent cancer cells were detached by incubating with Tryple Express Enzyme (1x; 12-604-013; Fisher Scientific) at 37°C for 5-10 minutes. APC conjugated anti-human MUC17 (BioLegend 395306) and CF640R conjugated PNA were diluted 1:200 in 2% FBS PBS and incubated with cells at 4°C for 1 hour for each stain. Cell-surface Tn-MUC1 was measured using anti-human Tn-MUC1 (FHD14210-100; ProteoGenix; 1:200), followed by Alexa Fluor 647-conjugated goat anti-human IgG (H+L) secondary antibody (Invitrogen; A56019; 1:200). MUC1 tandem-repeat epitopes were detected using anti-human MUC1 clone HMPV (BD Biosciences; 555925; 1:200), whereas the membrane-proximal MUC1-C epitope was detected using clone 6A6 (Absolute Antibody; Ab03830-1.1; 1:50), followed by Alexa Fluor 647-conjugated goat anti-mouse IgG (H+L) secondary antibody (Thermo Scientific; A-21235; 1:200). Antibodies were diluted in PBS containing 2% FBS at the indicated concentrations. Total cell-surface mucin expression level was assessed using biotinylated Mucin Probe StcE (Sigma-Aldrich; SAE0212) diluted 1:200 in FACS buffer. Bound probe was detected using streptavidin-Cy5 (Vector Laboratories; SA-1500-1; 1:200). Surface expression of membrane-tethered KLK5 was measured using anti-human KLK5 antibody (Novus Biologicals; NBP3-27961B; 1:50), followed by Alexa Fluor 647-conjugated goat anti-human IgG (H+L) secondary antibody (Invitrogen; A56019; 1:200). Unless otherwise indicated, cell-surface staining was performed for 1 h at 4 °C. Cells were washed and stained with DAPI containing 2% FBS PBS to assess cell viability before analyzing on a BD Fortessa X-20.

**Cytotoxicity assays**

For endpoint cytotoxicity assays, CBG luciferase-expressing target cells were cocultured with CAR-T cells or donor-matched untransduced T cells at the indicated effector-to-target (E:T) ratios in 200 µL of growth media specific to the target cell line, without IL-2, and cocultured in a 96-well plate for 30 hours at 37°C in 5% $CO_2$. Cells were then lysed using the Bright-Glo Luciferase Assay System (E2610; Promega), and luciferase activity was measured with a Biotek Neo2 luminescence plate reader. Specific lysis was calculated as [(luminescence target cell only) - (luminescence target cell + CAR-T cell)]/(luminescence target cell only) x 100%. CBG-eGFP-expressing target cells were seeded in flat-bottom 48- or 96-well plates (Corning) and allowed to adhere for at least 2 hours at 37°C in 5% $CO_2$. CAR-T cells or untransduced T cells were added in triplicate at various effector:target (E:T) ratios as indicated. CAR expression (%) was normalized using untransduced T cells for each donor. Plates were then incubated at 37°C for up to 6 days, with whole wells recorded every 2 hours using the IncuCyte Live Cell Analysis system. Tumor-cell growth and cytotoxicity were quantified from the total GFP-positive area using IncuCyte image-analysis software.

**Immunohistochemistry (IHC)**

After tumor-engrafted mice were euthanized, tumors were extracted and fixed in 4% PFA overnight, washed with PBS, and serial wash with 30%, 50%, 70% ethanol and stored in 70% EtOH until staining. Tissue slides were then made and embedded in paraffin. Slides were stained for CD3 and MUC17 (Novus Biologicals; NBP1-91013) by the specialized histopathology services core facility at MGH. All IHC slides were imaged on a Axio Scan.Z1 microscope and quantified the respective stains by QuPath (v0.5.1).

**Cell avidity measurement with acoustic force microscopy**

ASPC-1 and GSU cells were treated with KLK5 or the indicated enzyme under the conditions described above, washed, and seeded in poly-L-lysine-coated z-Movi chips (Lumicks) at a density of 60 x $10^6$ cells/mL and incubated for 2 hours. CAR-T cells were sorted using a Sony MAB900 (Sony Biotechnology Inc.) 48 hours before the avidity assessment, with anti-$(G_4S)_3$ linker conjugated with Alexa Fluor 647 and mCherry$^+$ gates to isolate only CAR+ T cells. Sorted CAR-T cells and untransduced T cells were stained with a CellTrace™ Far Red Proliferation Kit (Thermo Fisher Scientific). Each chip was first run with untransduced T cells to prevent nonspecific confounding binding of CAR-T cells, followed by CAR-T cells with three different bridges or HPA-based CAR, and finally untransduced T cells. To eliminate any potential bias in binding from earlier runs, the order of CAR-T cells to be tested was alternated. T cells were incubated for a 5-minute binding period and visualized on the z-Movi Cell Avidity Analyzer (Lumicks), while a gradient acoustic force of up to 1000 pN was applied. The percentage of cells bound as a function of the acoustic force applied was then analyzed using Ocean software (version 1.5.5, Lumicks). Each chip was run five times.

**RNA sequencing and gene-set enrichment analysis**

MUC17 CAR-T, MUC17 kCAR-T and AER-KLK5 MUC17 CAR-T cells were cocultured with GSU tumor cells at an E:T ratio of 2:1 for 48 h. CAR-T cells were generated from 6, 7 and 5 distinct healthy human donors for the MUC17 CAR, MUC17 kCAR and AER-KLK5 groups, respectively. Following coculture, T cells were collected in Zymo DNA/RNA Shield and submitted to Plasmidsaurus for RNA sequencing and primary bioinformatic analysis. Samples were processed using the Plasmidsaurus 3′-end-counting RNA-sequencing workflow with Illumina sequencing. Gene-level expression and differential expression analyses were performed using the standard Plasmidsaurus analysis pipeline. Gene-set enrichment analysis was performed using GSEApy with the MSigDB Hallmark gene sets.

**Statistical methods, sample sizes, data collection, and assumptions**

The sample sizes were selected on the basis of standards in the field and not pre-determined using statistical methods. For statistical comparisons, data distributions were assumed to be normal. Normality was tested for conditions with approximately ten or more data points. All the statistical analyses were performed using GraphPad Prism 8 software. All the experimental data are presented as mean ± s.d. or as box-and-whisker plots with the first and third quartiles (boxes), median and range of data, unless stated otherwise within figure legends. Appropriate statistical tests were used to analyze the data, as described in the figure legends.

### Supplementary Note 1

#### Thermodynamic model of CAR-antigen engagement across a mucin-rich glycocalyx

We constructed a thermodynamic model to examine how competition between CAR-antigen binding and glycocalyx-mediated repulsion could produce the non-monotonic relationship between mucin abundance and CAR engagement observed experimentally. The model was adapted from classical thermodynamic descriptions of receptor-mediated cell adhesion(*38*) and from a flexible polymer-brush model of the cancer glycocalyx(*39*). The model was intended to examine qualitative relationships between mucin abundance, CAR-antigen affinity, and CAR abundance rather than to quantitatively fit absolute molecular densities.

***Glycocalyx repulsion.*** We considered a CAR-T cell (cell 1) interacting with a mucin-bearing target cell (cell 2). The model includes CAR-antigen binding and nonspecific repulsion generated by the glycocalyx, while other receptor-ligand interactions were omitted.

The combined glycocalyx-density parameter within the cell-cell interface was defined as

$$C_G = C_{G1} + C_{G2}$$

Here, $C_{G1}$ and $C_{G2}$ represent the glycocalyx contributions from the CAR-T cell and target cell, respectively.

Following a flexible polymer-brush model, the repulsive free-energy density was written as

$$u_{\text{rep}}(S, C_G) = k_B T \left[ C_G + \frac{3S^2 C_G}{8N_K L_K^2} + \frac{2v_K C_G^2 N_K^2}{S} + \frac{Q_G^2 C_G^2}{2C_M S} \right]$$

Here, $S$ is the membrane separation, $N_K$ is the number of Kuhn segments per glycocalyx polymer, $L_K$ is the Kuhn length, $v_K$ is the excluded-volume parameter, $Q_G$ describes the effective polymer charge, and $C_M$ is the ionic-strength parameter.

The total repulsive free energy within the cell-cell contact area $A_c$ is therefore

$$G_{\text{rep}} = A_c u_{\text{rep}}(S, C_G)$$

***CAR-anbtigen binding.*** CAR-antigen interactions were treated as reversible elastic bonds. The separation-dependent two-dimensional binding constant was defined as

$$K(S) = K_L \exp\left[-\frac{k(S-L)^2}{2k_BT}\right]$$

where $K_L$ is the two-dimensional binding constant at the unstressed bond length, $L$ is the unstressed CAR-antigen bond length, and $k$ is the effective spring constant of the CAR-antigen bond.
At thermodynamic equilibrium, the CAR-antigen binding condition is

$$\frac{N_b}{A_c} = K(S)\left(\frac{N_1 - N_b}{A_1}\right)\left(\frac{N_2 - N_b}{A_2}\right)$$

where $N_1$ is the total number of CAR molecules on the CAR-T cell, $N_2$ is the total number of target antigens on the target cell, $N_b$ is the number of CAR-antigen bonds, and $A_1$ and $A_2$ are the cell-surface areas.

Variation of the total free energy with respect to contact area gives the area-balance condition

$$\frac{N_b}{A_c} = \frac{u_{\text{rep}}(S, C_G)}{k_BT}$$

Mechanical equilibrium in the direction normal to the cell surfaces gives

$$\frac{N_b}{A_c} k(S-L) + \frac{\partial u_{\text{rep}}}{\partial S} = 0$$

***Equilibrium membrane separation.*** Combining the area-balance and force-balance conditions yields a fifth-order polynomial in membrane separation, denoted S. In the numerical implementation, this equation was written as

$$t_1 S^5 + t_2 S^4 + C_G t_3 S^3 + C_G k t_4 S^2 + C_G^2 k L t_5 S + t_6 = 0,$$

with

$$t_1 = -\frac{3 C_G k}{8 N_K L_K^2}$$

$$t_2 = \frac{3 C_G k L}{8 N_K L_K^2}$$

$$t_3 = -k - \frac{3 k_B T}{4 N_K L_K^2}$$

$$t_4 = L - 2 v_K C_G N_K^2 - \frac{Q_G^2 C_G}{2 C_M}$$

$$t_5 = 2 v_K N_K^2 + \frac{Q_G^2}{2 C_M}$$

and

$$t_6 = 2 k_B T v_K C_G^2 N_K^2 + \frac{k_B T Q_G^2 C_G^2}{2 C_M}$$

The physically positive solution was determined numerically and defined as the equilibrium membrane separation, $S_{\mathrm{eq}}$.

***Calculation of equilibrium CAR engagement***

For convenience, the dimensionless repulsion term was defined as

$$R(S) = \frac{u_{\mathrm{rep}}(S, C_G)}{k_B T}$$

and

$$\zeta = \frac{A_1 A_2 R(S_{\mathrm{eq}})}{K(S_{\mathrm{eq}})}$$

Using the equilibrium separation, the number of CAR-antigen bonds was calculated as

$$N_b = \frac{1}{2}\left[N_1 + N_2 - \sqrt{(N_1 - N_2)^2 + 4\zeta}\right]$$

The corresponding equilibrium contact area was

$$A_c = \frac{N_b}{R(S_{\text{eq}})}$$

The primary model output was the fraction of CAR molecules engaged with antigen,

$$f_{\text{bound}} = \frac{N_b}{N_1}$$

Solutions producing negative values of $N_b$ or $A_c$ were assigned to the non-adherent state, with $N_b = 0$ and $A_c = 0$.

***Coupling mucin abundance to antigen abundance.*** To represent a mucin that contributes both to the glycocalyx barrier and to CAR-targeted antigen abundance, target-cell antigen abundance was coupled to the target-cell mucin-density parameter according to

$$N_2 = \alpha_{\text{mucin}} C_{G2} A_2$$

Thus, increasing $C_{G2}$ simultaneously increases the number of available CAR-binding sites and the repulsive contribution of the glycocalyx. $\alpha_{\text{mucin}}$ was used as a scaling parameter linking mucin abundance to antigen abundance and does not represent a measured stoichiometric number of epitopes per individual mucin molecule.

This coupling creates two competing effects of increasing mucin abundance: increasing antigen availability favors CAR-antigen binding, whereas increasing glycocalyx density increases the energetic cost of membrane apposition.

***Numerical implementation and parameter sweeps.*** The model was implemented in Julia. The fifth-order equilibrium equation was solved numerically using Roots.jl to obtain $S_{\text{eq}}$. $N_b$, $A_c$, and $f_{\text{bound}}$ were subsequently calculated from the equilibrium solution. For the simulations shown, the CAR-T-cell glycocalyx contribution $C_{G1}$ was held constant while $C_{G2}$ was varied. CAR-antigen affinity was varied using $K_L$ values of $10^{-8}$, $3 \times 10^{-8}$, $10^{-7}$, $3 \times 10^{-7}$, $10^{-6}$, $3 \times 10^{-6}$, and $10^{-5}$ $cm^2$. CAR abundance was varied using $N_1$ values of $1 \times 10^4$, $5 \times 10^4$, and $1 \times 10^5$ CAR molecules per cell. Unless otherwise indicated, all other parameters were held constant. The model therefore tests whether the experimentally observed non-monotonic dependence of CAR engagement on mucin abundance can arise from competition between increasing antigen availability and increasing glycocalyx-mediated repulsion. It was not used as a quantitative fit to absolute mucin or receptor densities. Simulations were performed using Julia 1.12.2 and Roots.jl version 2.2.10.

***Model assumptions.*** The model assumes thermodynamic equilibrium and does not explicitly include receptor-binding kinetics, intracellular signaling, active cytoskeletal forces, or time-dependent membrane deformation. CARs and antigens are treated as laterally well-mixed species, CAR-antigen bonds are represented by a common effective bond length and spring constant, and the glycocalyx is represented as a homogeneous flexible charged polymer layer. Spatial heterogeneity, receptor clustering, cooperative binding, and variation among individual mucin conformations are not explicitly represented. Because the mucin-density and mucin-to-antigen coupling parameters are scaled model quantities, the simulations are interpreted qualitatively rather than as absolute molecular-density predictions.

**Supplementary Figures 1-6**

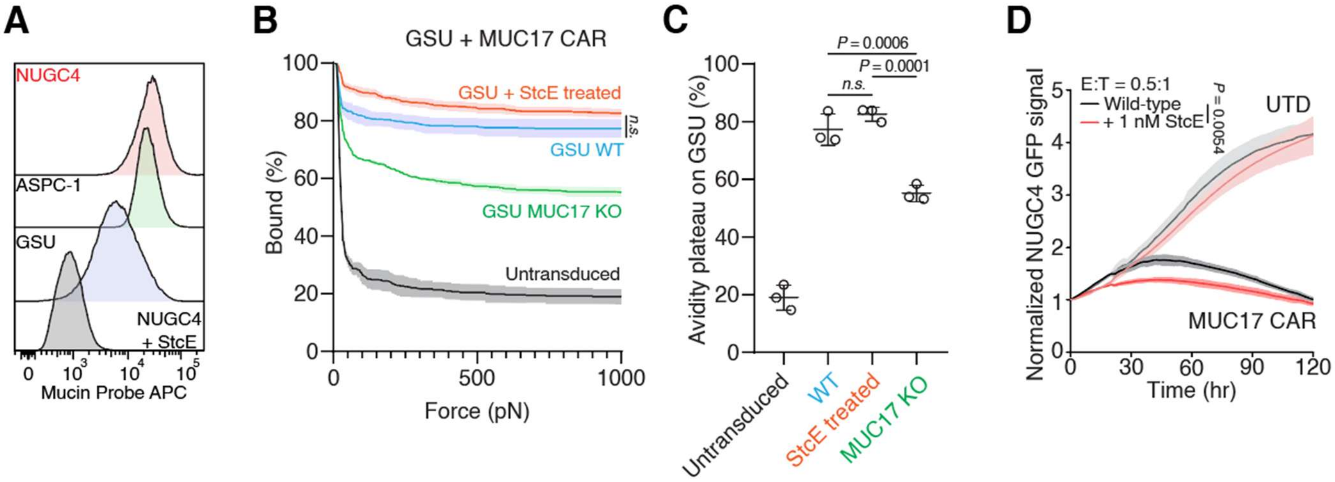


**Supplementary Fig 1. StcE-mediated glycocalyx remodeling enhances MUC17 CAR-T-cell engagement.** (**A**) Flow cytometry analysis of cell-surface mucin expression across NUGC4, ASPC-1, and GSU. (**B** and **C**) Interaction strength between untreated or StcE-treated NUGC4 cells, MUC17-knockout GSU cells and either MUC17 CAR-T or untransduced T cells. The percentage of T cells remaining bound to target cells is shown as the applied acoustic force is increased from 0 to 1,000 pN (**B**). The percentage of CAR-T cells remaining bound at the plateau of an avidity at 1,000 pN is also quantified (**C**. Results are mean ± s.e.m. for force-ramp measurements and mean ± s.d. for avidity-plateau measurements from n = 3 independent measurements. (**D**) Representative real-time cytotoxicity assay of MUC17 CAR-T cells against untreated or NUGC4 cells at 1:1 E:T ratio with 1nM StcE. Results are mean ± s.d. of n = 3 independent measurements. In (**C**), statistical analysis was performed by one-way ANOVA with Tukey's post hoc tests. In (**B**) and (**D**), statistical analysis was performed by two-way ANOVA with correction for multiple comparisons.

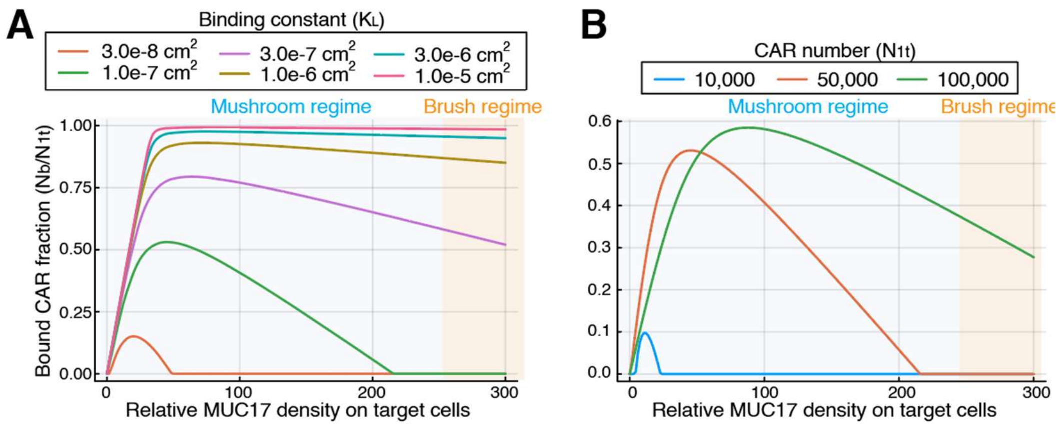


**Supplementary Fig. 2. Theoretical modeling of mucin density-dependent CAR-T-cell engagement.** (**A** amd **B**) Theoretical predictions of the relationship between mucin density and MUC17 CAR-T-cell activity as a function of CAR-antigen binding affinity (**a**) or CAR surface expression (**b**).

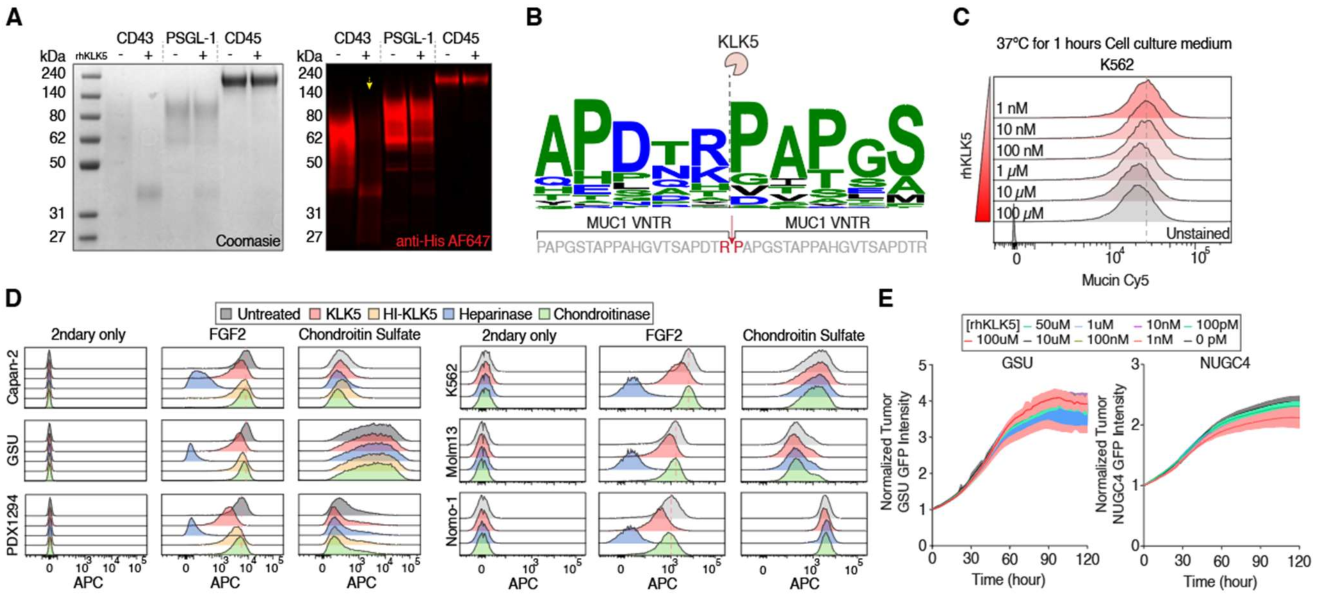


**Supplementary Fig. 3. Biochemical and cellular characterization of KLK5-mediated glycocalyx remodeling.** (**A**) Coomassie staining and western blot analysis of recombinant CD43, PSGL-1 and CD45 following treatment with KLK5. (**B**) LC-MS/MS glycopeptide analysis of MUC1 following KLK5 treatment. (**C**) Flow cytometry analysis of cell-surface mucins in K562 after the indicated concentration of recombinant human KLK5 (rhKLK5). (**D**) Flow cytometry analysis of FGF2 binding and chondroitin sulfate on the indicated cell lines after treatment with KLK5, heat-inactivated KLK5, heparinase, chondroitinase. (**E**) Representative real-time tumor growth assay of GSU and NUGC4 cells with treatment of the indicated rhKLK5. Results are mean ± s.d. of n = 3 independent measurements.

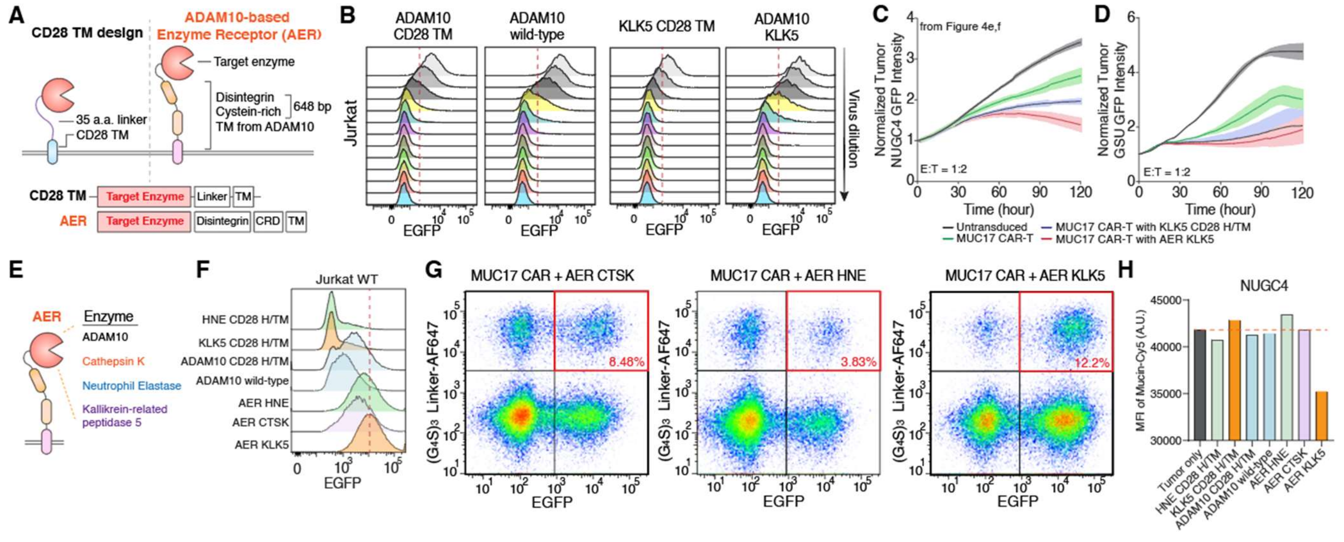

**Supplementary Fig. 4. Optimization and characterization of membrane-tethered enzyme receptor designs.** (**A**) Schematics illustrating membrane-tethered enzyme strategies using a CD28 transmembrane (TM)-based design and an engineered ADAM10-based enzyme receptor (AER). (**B**) Flow cytometry analysis of construct expression in Jurkat cells transduced with ADAM10-CD28 TM, wild-type ADAM10, KLK5-CD28 TM or AER-KLK5 constructs. Each row represents a 3-fold serial dilution of lentiviral input. (**C**) Schematic of the AER architecture incorporating the indicated protease domains: ADAM10, cathepsin K, human neutrophil elastase or KLK5. (**D**) Flow cytometry analysis of expression of the indicated AER or CD28-based constructs in wild-type Jurkat cells. (**E**) Representative flow cytometry analysis of G4S and GFP expression in primary human T cells transduced with the indicated constructs. The boxed population indicates $G4S^{+}GFP^{+}$ double-positive cells. (**F**) Quantification of cell-surface mucin levels on NUGC4 cells following 24 h coculture at 37 °C with T cells expressing the indicated constructs. Mucin levels are shown as mean fluorescence intensity (MFI). (**G** and **H**) Representative real-time cytotoxicity assays of untransduced T cells, MUC17 CAR-T cells, MUC17 CAR-T cells expressing CD28 TM-tethered KLK5, and MUC17 CAR-T cells expressing AER-KLK5 against NUGC4 (**G**) and GSU (**H**) target cells at an E:T ratio of 1:2. Tumor-cell growth was normalized to the tumor-cell signal at day 0. Data are mean ± s.d. from n = 3 distinct human blood-cell donors. Data for untransduced T cells, conventional MUC17 CAR-T cells and AER-KLK5 MUC17 CAR-T cells are the same datasets shown in (**Fig. 4E and F**); the CD28 TM-tethered KLK5 group is shown here for comparison.

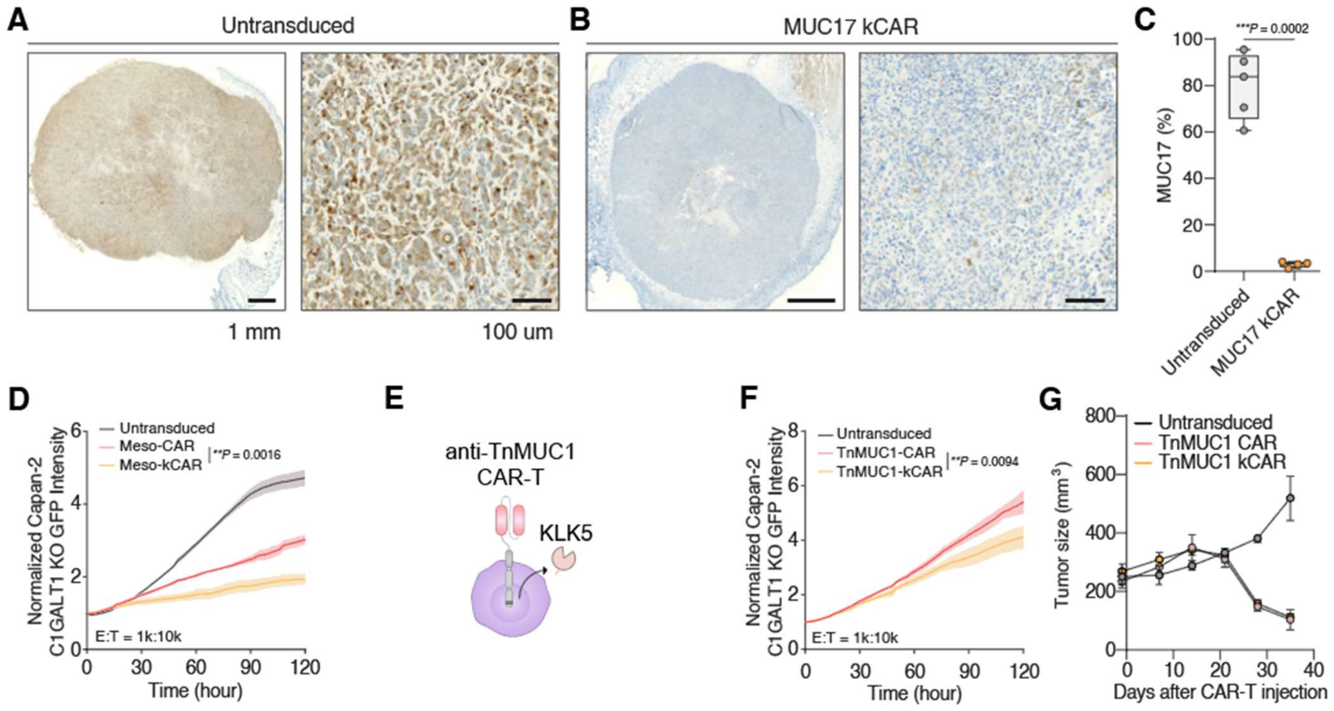


**Supplementary Fig. 5. Characterization of KLK5-secreting CAR-T cells across distinct target antigens and tumor models.** (**A** and **B**) Representative immunohistochemistry (IHC) images of human MUC17 expression in formalin-fixed, paraffin-embedded ASPC-1 tumors from mice treated with untransduced T cells (**A**) or MUC17 kCAR-T cells (**B**) in the experiment shown in (**Fig. 5J and K)**. Scale bars, 1 mm (left) and 100 µm (right). (**C**) Quantification of MUC17-positive cells in tumors from **a,b**. Boxes and whiskers indicate the first and third quartiles (boxes), median and range from n = 4 tumors per group available for analysis; mice that died before tissue collection were excluded. (**D** and **E**) Representative real-time cytotoxicity assays of mesothelin CAR-T and mesothelin kCAR-T cells (**D**) and Tn-MUC1 CAR-T and Tn-MUC1 kCAR-T cells (**E**) against Capan-2 *C1GALT1* KO cells with 1:10 E:T ratio. Results are mean ± s.d. of n = 3 independent measurements. (**F**) *in vivo* antitumor activity of TnMUC1 kCAR-T cells in the Capan-2 *C1GALT1* KO xenograft model corresponding to (**Fig. 5L)**. NSG mice were injected subcutaneously with $5 \times 10^6$ Capan-2 *C1GALT1* KO cells on day -14. On day 0, mice received $2 \times 10^6$ untransduced T cells, TnMUC1 CAR-T cells or TnMUC1 kCAR-T cells (n = 5 mice per group), and tumor growth was monitored over time. Data are mean ± s.e.m. In (**C**), statistical analysis was performed by two-tailed t-tests. In (**D** and **E**) statistical analysis was performed by two-way ANOVA with correction for multiple comparisons.

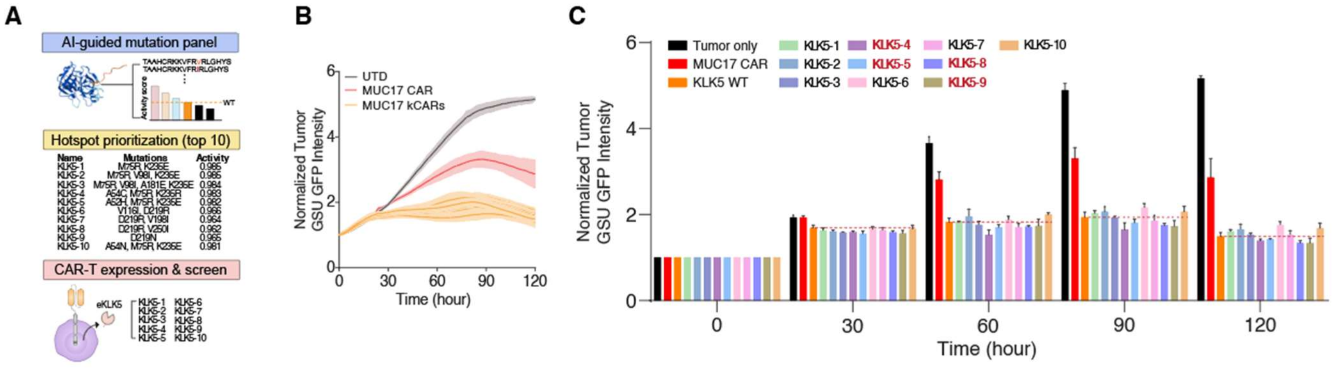


**Supplementary Fig. 6. Functional screening of AI-guided KLK5 variants in CAR-T cells.** (**A**) Schematic of the AI-guided KLK5 engineering workflow. Candidate residues were prioritized based on predicted mutational effects, and the top-ranked KLK5 variants were selected for expression and functional screening in CAR-T cells. (**B**) Representative real-time real-time cytotoxicity assay of untransduced T cells (UTD), conventional MUC17 CAR-T cells and MUC17 CAR-T cells expressing engineered KLK5 variants against GSU target cells at 1:5 E:T ratio. Results are mean ± s.d. of n = 3 independent measurements. (**C**) Comparison of normalized GSU tumor GFP intensity from the same dataset shown in **(B)** at 0, 30, 60, 90 and 120 h for wild-type KLK5 and the indicated KLK5 variants. Data are mean ± s.d. of n = 3 independent measurements.

**Supplemental Table 1. Complete sequence for MUC17 CAR secreting enzymes and membrane-tethered enzymes**

| | |
|---|---|
| MUC17 CAR with KLK5-His<br><br>MUC17 CAR<br><br>KLK5 | MALPVTALLLPLALLLHAARPQVQLQQWGAGLLKPSETLSLTCAVYGGSFSGY<br>YWSWIRQPPGKCLEWIGDIDASGSTKYNPSLKSRVTISLDTSKNQFSLKLNSVTA<br>ADTAVYFCARKKYSTVWSYFDNWGQGTLVTVSSGGGGSGGGGSGGGGSSYEL<br>TQPSSVSVPPGQTASITCSGDKLGDKYASWYQQKPGQSPVLVIYQDRKRPSGVP<br>ERFSGSNSGNTATLTISGTQAMDEADYYCQAWGSSTAVFGCGTKLTVLTTTPAPR<br>PPTPAPTIASQPLSLRPEACRPAAGGAVHTRGLDFACDIYIWAPLAGTCGVLLLSL<br>VITLYCKRGRKKLLYIFKQPFMRPVQTTQEEDGCSCRFPEEEEGGCELRVKFSRS<br>ADAPAYQQGQNQLYNELNLGRREEYDVLDKRRGRDPEMGGKPRRKNPQEGLY<br>NELQKDKMAEAYSEIGMKGERRRGKGHDGLYQGLSTATKDTYDALHMQALPP<br>RSGGGGEGRGSLLTCGDVEENPGPRMATARPPWMWVLCALITALLLGVTEHVL<br>ANNDVSCDHPSNTVPSGSNQDLGAGAGEDARSDDSSSRIINGSDCDMHTQPWQ<br>AALLLRPNQLYCGAVLVHPQWLLTAAHCRKKVFRVRLGHYSLSPVYESGQQMF<br>QGVKSIPHPGYSHPGHSNDLMLIKLNRRIRPTKDVRPINVSSHCPSAGTKCLVSG<br>WGTTKSPQVHFPKVLQCLNISVLSQKRCEDAYPRQIDDTMFCAGDKAGRDSCQ<br>GDSGGPVVCNGSLQGLVSWGDYPCARPNRPGVYTNLCKFTKWIQETIQANSHH<br>HHHH* |
| MUC17 with KLK-5-Linker-Myc-CD28 TM<br><br>MUC17 CAR<br><br>KLK5<br><br>CD28 TM | MALPVTALLLPLALLLHAARPQVQLQQWGAGLLKPSETLSLTCAVYGGSFSGY<br>YWSWIRQPPGKCLEWIGDIDASGSTKYNPSLKSRVTISLDTSKNQFSLKLNSVTA<br>ADTAVYFCARKKYSTVWSYFDNWGQGTLVTVSSGGGGSGGGGSGGGGSSYEL<br>TQPSSVSVPPGQTASITCSGDKLGDKYASWYQQKPGQSPVLVIYQDRKRPSGVP<br>ERFSGSNSGNTATLTISGTQAMDEADYYCQAWGSSTAVFGCGTKLTVLTTTPAPR<br>PPTPAPTIASQPLSLRPEACRPAAGGAVHTRGLDFACDIYIWAPLAGTCGVLLLSL<br>VITLYCKRGRKKLLYIFKQPFMRPVQTTQEEDGCSCRFPEEEEGGCELRVKFSRS<br>ADAPAYQQGQNQLYNELNLGRREEYDVLDKRRGRDPEMGGKPRRKNPQEGLY<br>NELQKDKMAEAYSEIGMKGERRRGKGHDGLYQGLSTATKDTYDALHMQALPP<br>RSGGGGEGRGSLLTCGDVEENPGPRMATARPPWMWVLCALITALLLGVTEHVL<br>ANNDVSCDHPSNTVPSGSNQDLGAGAGEDARSDDSSSRIINGSDCDMHTQPWQ<br>AALLLRPNQLYCGAVLVHPQWLLTAAHCRKKVFRVRLGHYSLSPVYESGQQMF<br>QGVKSIPHPGYSHPGHSNDLMLIKLNRRIRPTKDVRPINVSSHCPSAGTKCLVSG<br>WGTTKSPQVHFPKVLQCLNISVLSQKRCEDAYPRQIDDTMFCAGDKAGRDSCQ<br>GDSGGPVVCNGSLQGLVSWGDYPCARPNRPGVYTNLCKFTKWIQETIQANS<br>GGGGSGGGGSGGGGSGGGGSGGGGSGGGGSGGGGSEQKLISEEDLIEVMYPPP<br>YLDNEKSNGTIIHVKGKHLCPSPLFPGPSKPFWVLVVVGGVLACYSLLVTVAFIIF<br>WV* |
| MUC17 CAR with AER-KLK5<br><br>MUC17 CAR | MALPVTALLLPLALLLHAARPQVQLQQWGAGLLKPSETLSLTCAVYGGSFSGY<br>YWSWIRQPPGKCLEWIGDIDASGSTKYNPSLKSRVTISLDTSKNQFSLKLNSVTA<br>ADTAVYFCARKKYSTVWSYFDNWGQGTLVTVSSGGGGSGGGGSGGGGSSYEL<br>TQPSSVSVPPGQTASITCSGDKLGDKYASWYQQKPGQSPVLVIYQDRKRPSGVP<br>ERFSGSNSGNTATLTISGTQAMDEADYYCQAWGSSTAVFGCGTKLTVLTTTPAPR<br>PPTPAPTIASQPLSLRPEACRPAAGGAVHTRGLDFACDIYIWAPLAGTCGVLLLSL<br>VITLYCKRGRKKLLYIFKQPFMRPVQTTQEEDGCSCRFPEEEEGGCELRVKFSRS<br>ADAPAYQQGQNQLYNELNLGRREEYDVLDKRRGRDPEMGGKPRRKNPQEGLY |

| | |
|---|---|
| ADAM10<br>SP<br><br>KLK5<br><br>Disintegrin<br><br>Cystein<br>Rich<br><br>TM | NELQKDKMAEAYSEIGMKGERRRGKGHDGLYQGLSTATKDTYDALHMQALPP<br>RSGGGGEGRGSLLTCGDVEENPGPRMVLLRVLILLLSWAAGMGGVTEHVLANN<br>DVSCDHPSNTVPSGSNQDLGAGAGEDARSDDSSSRIINGSDCDMHTQPWQAAL<br>LLRPNQLYCGAVLVHPQWLLTAAHCRKKVFRVRLGHYSLSPVYESGQQMFQGV<br>KSIPHPGYSHPGHSNDLMLIKLNRRIRPTKDVRPINVSSHCPSAGTKCLVSGWGT<br>TKSPQVHFPKVLQCLNISVLSQKRCEDAYPRQIDDTMFCAGDKAGRDSCQGDS<br>GGPVVCNGSLQGLVSWGDYPCARPNRPGVYTNLCKFTKWIQETIQANSQPICG<br>NGMVEQGEECDCGYSDQCKDECCFDANQPEGRKCKLKPGKQCSPSQGPCCTA<br>QCAFKSKSEKCRDDSDCAREGICNGFTALCPASDPKPNFTDCNRHTQVCINGQC<br>AGSICEKYGLEECTCASSDGKDDKELCHVCCMKKMDPSTCASTGSVQWSRHFS<br>GRTITLQPGSPCNDFRGYCDVFMRCRLVDADGPLARLKKAIFSPELYENIAEWIV<br>AHWWAVLLMGIALIMLMA* |
| KLK5-35<br>aa GGGS<br>Linker-<br>TnMUC1<br><br>KLK5<br><br>Linker<br><br>TnMUC1 | VTEHVLANNDVSCDHPSNTVPSGSNQDLGAGAGEDARSDDSSSRIINGSDCDM<br>HTQPWQAALLLRPNQLYCGAVLVHPQWLLTAAHCRKKVFRVRLGHYSLSPVYE<br>SGQQMFQGVKSIPHPGYSHPGHSNDLMLIKLNRRIRPTKDVRPINVSSHCPSAGT<br>KCLVSGWGTTKSPQVHFPKVLQCLNISVLSQKRCEDAYPRQIDDTMFCAGDKA<br>GRDSCQGDSGGPVVCNGSLQGLVSWGDYPCARPNRPGVYTNLCKFTKWIQETI<br>QANSGGGGSGGGGSGGGGSGGGGSGGGGSGGGGSGGGGSQVQLQQSDAELV<br>KPGSSVKISCKASGYTFTDHAIHWVKQKPEQGLEWIGHFSPGNTDIKYNDKFKG<br>KATLTVDRSSSTAYMQLNSLTSEDSAVYFCKTSTFFFDYWGQGTTLTVSSGGGGS<br>GGGGSGGGGSELVMTQSPSSLTVTAGEKVTMICKSSQSLLNSGDQKNYLTWYQ<br>QKPGQPPKLLIFWASTRESGVPDRFTGSGSGTDFTLTISSVQAEDLAVYYCQNDY<br>SYPLTFGAGTKLELKHHHHHH* |
| KLK5-35<br>aa GGGS<br>Linker-<br>CD19<br><br>KLK5<br><br>Linker<br><br>CD19 | VTEHVLANNDVSCDHPSNTVPSGSNQDLGAGAGEDARSDDSSSRIINGSDCDM<br>HTQPWQAALLLRPNQLYCGAVLVHPQWLLTAAHCRKKVFRVRLGHYSLSPVYE<br>SGQQMFQGVKSIPHPGYSHPGHSNDLMLIKLNRRIRPTKDVRPINVSSHCPSAGT<br>KCLVSGWGTTKSPQVHFPKVLQCLNISVLSQKRCEDAYPRQIDDTMFCAGDKA<br>GRDSCQGDSGGPVVCNGSLQGLVSWGDYPCARPNRPGVYTNLCKFTKWIQETI<br>QANSGGGGSGGGGSGGGGSGGGGSGGGGSGGGGSGGGGSEIVMTQSPATLSLS<br>PGERATLSCRASQDISKYLNWYQQKPGQAPRLLIYHTSRLHSGIPARFSGSGSGT<br>DYTLTISSLQPEDFAVYFCQQGNTLPYTFGQGTKLEIKGGGGSGGGGSGGGGSG<br>GGGSQVQLQESGPGLVKPSETLSLTCTVSGVSLPDYGVSWIRQPPGKGLEWIGVI<br>WGSETTYYQSSLKSRVTISKDNSKNQVSLKLSSVTAADTAVYYCAKHYYYGGS<br>YAMDYWGQGTLVTVSSHHHHHH* |
| KLK5-35<br>aa GGGS<br>Linker-<br>Mesothelin<br><br>KLK5<br><br>Linker<br><br>CD19 | VTEHVLANNDVSCDHPSNTVPSGSNQDLGAGAGEDARSDDSSSRIINGSDCDM<br>HTQPWQAALLLRPNQLYCGAVLVHPQWLLTAAHCRKKVFRVRLGHYSLSPVYE<br>SGQQMFQGVKSIPHPGYSHPGHSNDLMLIKLNRRIRPTKDVRPINVSSHCPSAGT<br>KCLVSGWGTTKSPQVHFPKVLQCLNISVLSQKRCEDAYPRQIDDTMFCAGDKA<br>GRDSCQGDSGGPVVCNGSLQGLVSWGDYPCARPNRPGVYTNLCKFTKWIQETI<br>QANSGGGGSGGGGSGGGGSGGGGSGGGGSGGGGSGGGGSQVQLQQSGPELEK<br>PGASVKISCKASGYSFTGYTMNWVKQSHGKSLEWIGLITPYNGASSYNQKFRG<br>KATLTVDKSSSTAYMDLLSLTSEDSAVYFCARGGYDGRGFDYWGQGTTVTVSS<br>GGGGSGGGGSGGGGSDIELTQSPAIMSASPGEKVTMTCSASSSVSYMHWYQQK<br>SGTSPKRWIYDTSKLASGVPGRFSGSGSGNSYSLTISSVEAEDDATYYCQQWSG<br>YPLTFGAGTKLEIHHHHHH* |
